# Methylation and Gene Expression Differences Between Reproductive Castes of Bumblebee Workers

**DOI:** 10.1101/517698

**Authors:** Hollie Marshall, Zoë N. Lonsdale, Eamonn B. Mallon

## Abstract

Phenotypic plasticity is the production of multiple phenotypes from a single genome and is notably observed in social insects. Multiple epigenetic mechanisms have been associated with social insect plasticity, with DNA methylation being explored to the greatest extent. DNA methylation is thought to play a role in caste determination in Apis mellifera, and other social insects, but there is limited knowledge on it’s role in other bee species. In this study we analysed whole genome bisulfite sequencing and RNA-seq data sets from head tissue of reproductive and sterile castes of the eusocial bumblebee Bombus terrestris. We found genome-wide methylation in B. terrestris is similar to other social insects and does not differ between reproductive castes. We did, however, find differentially methylated genes between castes, which are enriched for multiple biological processes including reproduction. However we found no relationship between differential methylation and differential gene expression or differential exon usage between castes. Our results also indicate high inter-colony variation in methylation. These findings suggest methylation is associated with caste differences but may serve an alternate function, other than direct caste determination in this species. This study provides the first insights into the nature of a bumblebee caste specific methylome as well as it’s interaction with gene expression and caste specific alternative splicing, providing greater understanding of the role of methylation in phenotypic plasticity within social bee species. Future experimental work is needed to determine the function of methylation and other epigenetic mechanisms in social insects.

**Impact Summary:** Social insects, such as ants, termites, bees and wasps, can produce individuals with extreme physical and behavioural differences within the same colony known as castes (e.g. workers/soldiers/queens). These individuals have similar genomes and many studies have associated epigenetic mechanisms with the differences observed. Epigenetic modifications are changes that affect how genes are expressed without changing the underlying DNA code. Here we investigated differences in DNA methylation (a well researched modified base) between different reproductive castes of the bumblebee, *Bombus terrestris*, an economically and environmentally important pollinator species. We found *B. terrestris* has a similar methylation profile to other social insect species in terms of the distribution of methylation throughout the genome and the relationship between methylation and gene expression. Genes that have differences in methylation between reproductive castes are involved in multiple biological processes, including reproduction, suggesting methylation may hold multiple functions in this species. These differentially methylated genes are also different to differentially methylated genes identified between honeybee reproductive castes, again suggesting methylation may have a variable function. These findings provide greater understanding of the role of methylation in caste determination in social insect species.

## Introduction

Phenotypic plasticity is the production of multiple phenotypes from a single genome. It plays a crucial role in the adaptive capabilities of species (Chevin *et al.*, 2010) and is notably observed in social insects. Social insects exhibit, sometimes extreme, morphological and behavioural differences within a single colony known as castes. The mechanisms by which species develop differences between castes are diverse; some species use only environmental queues whilst others rely only on inherited changes, with many species falling somewhere in between these two extremes (Matsuura *et al.*, 2018). For example some ant species from the *Pogonomyrmex* genus have purely genetic caste determination (Mott *et al.*, 2015). On the other hand, many ant species undergo caste determination in response to only the environment, indicating their genomes must contain the code for all caste possibilities, with the phenotype potentially determined by epigenetic factors (Bonasio *et al.*, 2012).

Multiple epigenetic mechanisms have been associated with social insect plasticity. Histone modifications have been shown to be involved with plasticity, for example changes in histone acetylation alter the behaviour of major workers of the ant species *Camponotus floridanus*, making them more similar to the behaviour of minor workers (Simola *et al.*, 2016). Variation in microRNA expression levels has been identified in both honeybee (Ashby *et al.*, 2016) and bumblebee (Collins *et al.*, 2017) castes. However the most active research in this area has been focused on DNA methylation (Glastad *et al.*, 2015).

DNA methylation is the addition of a methyl group to a cytosine nucleotide. In mammals, methylation primarily occurs in a CpG context (CpG referring to a cytosine base immediately followed by a guanine base), the percentage of CpG’s methylated is usually over 70%, with methylation serving to repress gene expression when occurring in promoter regions (Feng *et al.*, 2010). However in insects, it is generally found in much lower quantities, ranging from zero methylation in most Diptera species studied, to >2% in Hymenoptera and up to 14% in some species of Blattodea (Provataris *et al.*, 2018). It is also enriched in gene bodies rather than throughout the genome, as in mammals (Fang *et al.*, 2012; Wang *et al.*, 2013), with a possible role in alternative splicing (Bonasio *et al.*, 2012).

DNA methylation has been associated with the switching of worker castes in honeybees (Herb *et al.*, 2012). A major finding was that silencing of the *Dnmt3* gene (involved in methylation establishment) in larvae produced queens rather than workers (Kucharski *et al.*, 2008). DNA methylation has also been linked with alternative splicing differences between castes in two ant species (Bonasio *et al.*, 2012) and is thought to be involved in caste determination in *Copidosoma koehleri*, a species of primitively social wasp (Shaham *et al.*, 2016).

However, it is clear DNA methylation is not a conserved mechanism in Hymenoptera for caste differentiation. No association between caste and methylation has been found in a number of wasp and ant species (Standage *et al.*, 2016; Patalano *et al.*, 2015). A recent knockdown of *Dnmt1* in the red flour bettle (*Tribolium castaneum*), a species with no apparent DNA methylation, caused developmental arrest in embryos, questioning the role of *Dnmt* enzymes in some insects. Additionally, the statistical methods of previous next generation sequencing analyses on social insect methylation have been brought into question (Libbrecht *et al.*, 2016).

A greater variety of species are needed to begin to understand the role of DNA methylation in social insect caste determination. Here, we assess whole genome methylation differences between reproductive castes of the bumblebee, *Bombus terrestris*, with an aim to investigate the role of methylation in caste determination in this species. Bumblebees are primitively eusocial and are an important pollinator species, both economically and environmentally. They are generalist pollinators and are keystone species in some ecosystems (Woodard *et al.*, 2015). *B. terrestris* colonies are annual and are founded by a singly-mated queen in early spring, she will lay diploid eggs resulting in female workers and later switch to male haploid eggs, known as the switching point (Bloch, 1999). A competition phase then occurs between queens and workers, where some workers will become reproductive and produce their own haploid sons (Alaux *et al.*, 2006), this results in distinct reproductive worker castes within the colony. Multiple recent studies have highlighted *B. terrestris* as an ideal organism to assess methylation as a potential regulatory mechanism for reproductive caste determination (Li *et al.*, 2018; Lonsdale *et al.*, 2017; Amarasinghe *et al.*, 2014).

Methylation regulatory genes were identified in the bumblebee genome and have since been shown to have varying expression levels between queens, workers and drones (Li *et al.*, 2018). Additionally, genes showing allele-specific methylation and gene expression have been identified and are enriched in reproductive related processes (Lonsdale *et al.*, 2017). Most importantly, experimental changes in methylation in *B. terrestris* workers has been shown to alter levels of reproductive behaviour (Amarasinghe *et al.*, 2014). Whilst these studies highlight differences in methylation between *B. terrestris* castes it is still unclear where those differences are within the genome and also whether methylation differences are related to changes in gene expression, potentially leading to caste differentiation.

In this study, we compared whole genome bisulfite sequencing datasets from reproductive and sterile worker castes of *B. terrestris*, allowing us to identify differences in methylation at base-pair resolution throughout the genome. We then linked these data with gene expression data for the same individuals to identify a potential relationship between gene expression and methylation regarding reproductive caste determination. We hypothesise genome methylation in *B. terrestris* will show a similar profile to other social insects; with enrichment in coding regions and with more highly methylated genes being associated with more highly expressed genes (Patalano *et al.*, 2015; Glastad *et al.*, 2016; Bonasio *et al.*, 2012). If methylation plays a role in caste determination we would expect to find differentially methylated genes between castes, with functions related to reproduction. We would also expect any differentially methylated genes between castes to be enriched for differentially expressed genes or genes which have different exon usage between castes. Additionally if there is a conserved role for methylation in caste determination in Hymenoptera we would expect to find orthologous genes differentially methylated between *B. terrestris* reproductive castes and *A. mellifera* reproductive castes.

## Methods

### Bee husbandry and tissue sampling

Three *B. terrestris* colonies, from Agralan, UK, were reared in constant red light at 26°C and 60% humidity. They were fed 50% v/v apiary solution (Meliose-Roquette, France) and pollen (Percie du set, France) *ad libitum*. Callow workers, less than 24 hours old, were taken from each colony and placed in small rearing boxes of five individuals.

The worker bees were sacrificed at six days old. For each bee, the head was snap frozen in liquid nitrogen. Through dissection in 1% PBS solution, the reproductive status of each bee was determined and classed as either reproductive, sterile, or intermediate. Workers were classed as having developed ovaries, and therefore reproductive, if the largest oocyte was larger than the trophocyte follicle (Duchateau and Velthuis, 1988). This measurement is tightly correlated with reproductive status (Geva *et al.*, 2005; Foster *et al.*, 2004). The ovaries of each worker were weighed, and the length of the largest oocyte was measured using ImageJ v.1.50e (Schneider *et al.*, 2012) (supplementary 1.0.0). Worker ‘reproductiveness’ was classified on a scale from 0-4 based on Duchateau and Velthuis (1988), 0 begin completely sterile (Fig.1a) and 4 having fully developed ovaries (Fig.1b).

**Figure 1:**
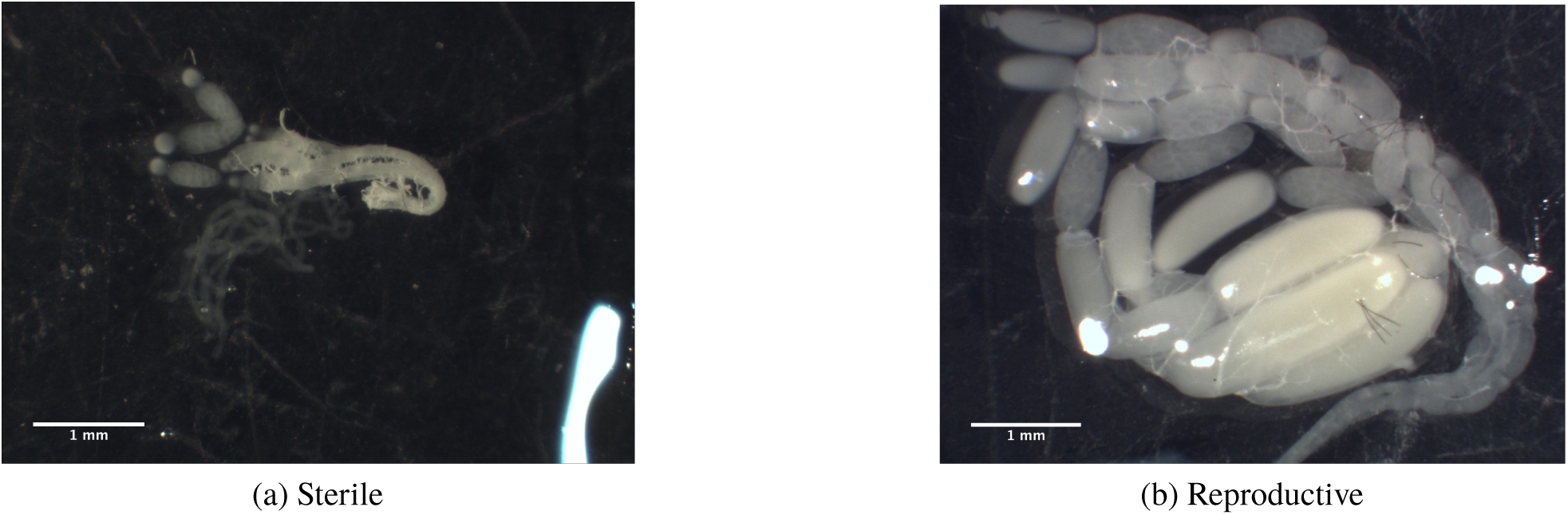
(a) one half of a pair of ovaries from a sterile bumblebee worker, with a score of 0. (b) one half of a pair of ovaries from a reproductive bumblebee worker, with a score of 4. Scores generated following Duchateau and Velthuis (1988).

### RNA and DNA extraction and sequencing

Three reproductive individuals and three sterile individuals from each of the three colonies were selected for RNA and DNA extraction, Fig.2. Heads were cut in half (using a lateral incision central between the eyes). Each head half was randomly allocated for either DNA/RNA extraction to avoid left/right hemisphere bias. RNA was extracted using the Sigma-Aldrich GenElute Mammalian Total RNA Miniprep kit and DNA was extracted using the Qiagen DNeasy blood and Tissue kit, individually for each half head per sample, following manufacturers protocols. The extracted RNA was treated with DNase and the extracted DNA was treated with RNase. DNA was pooled per colony and reproductive status, i.e. the three reproductive samples from a single colony were pooled to create one representative reproductive sample for that colony, Fig.2. RNA samples were processed individually. DNA and RNA quality and quantity was determined by Nanodrop and Qubit® fluorometers (supplementary 1.0.1 and 1.0.2). A total of 18 RNA samples (three individuals per reproductive status for each of the three colonies) were sent for 100bp paired-end sequencing and six pooled DNA samples (one sample per reproductive status per colony consisting of three individuals per pool) were sent for 100bp paired-end bisulfite sequencing on a HiSeq 2000 machine (Illumina, Inc.) by BGI Tech Solution Co., Ltd.(Hong Kong). Library preparation was carried out by BGI. A 1% lambda spike was included as an unmethylated control in each whole genome bisulfite sequencing (WGBS) library. Directional bisulfite libraries were prepared by fragmenting genomic DNA to 100-300bp by sonication, DNA-end repair was then carried out along with ligation of methylated sequencing adaptors. The ZYMO EZ DNA Methylation-Gold kit was used for bisulfite treatment with subsequent desaltation, size selection, PCR amplification and final size selection before sequencing. RNA-seq libraries were prepared by using magnetic beads with an Oligo (dT) to enrich for mRNA, the mRNA was then fragmented and cDNA was synthesised using random hexamer priming, size selection and PCR amplification were then carried out prior to sequencing.

**Figure 2:**
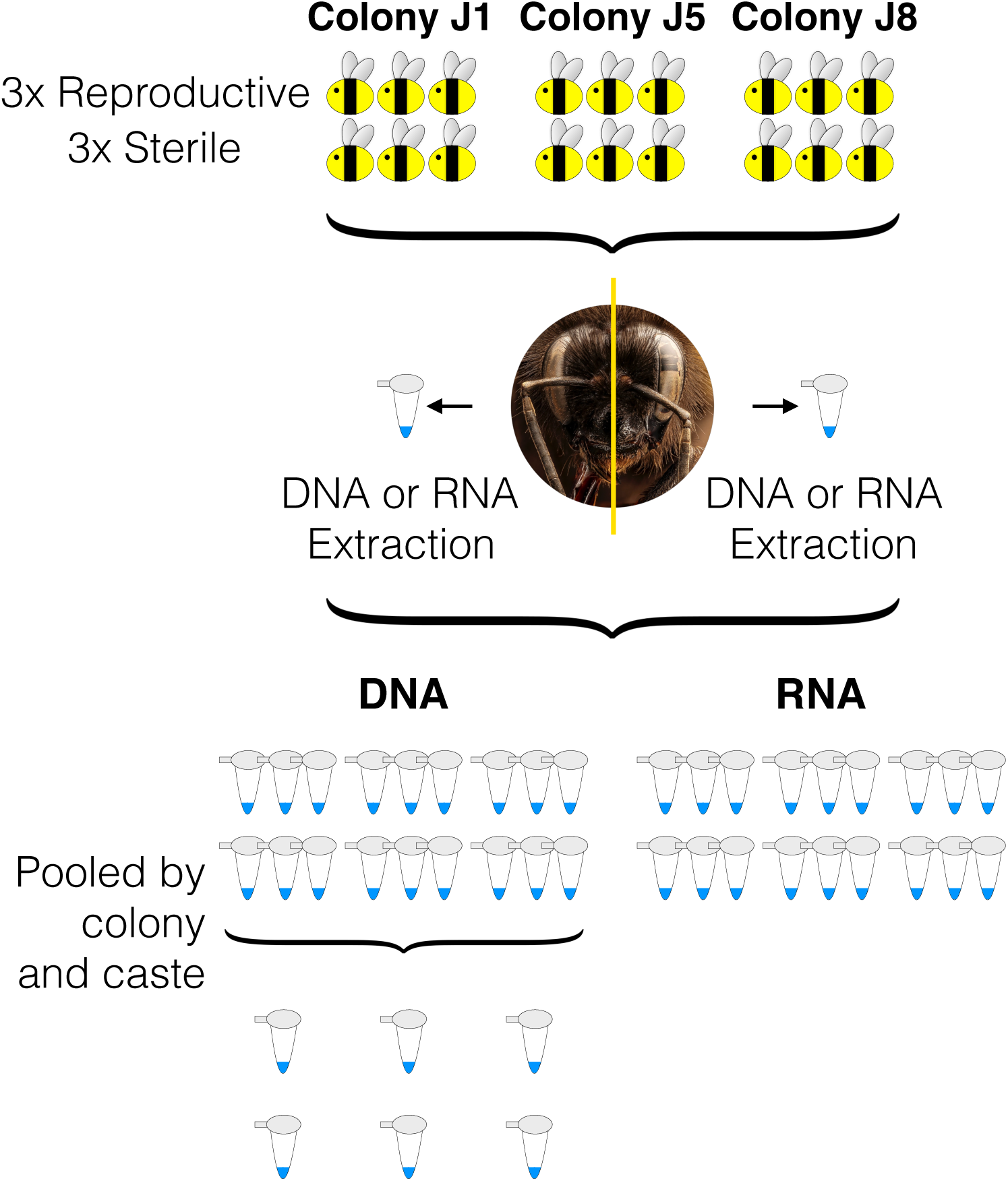
Overview of sample preparation for sequencing. Three reproductive workers and three sterile workers were selected from three colonies (J1, J5 and J8 represent colony names). Half of each head was randomly allocated for RNA/DNA extraction. All 18 RNA samples were sent for RNA-Seq (three of each caste from each colony). DNA samples were pooled by colony and caste creating a representative reproductive and sterile samples per colony.

### Differential expression and alternative splicing

Low quality bases were removed from the RNA-Seq libraries using CutAdapt v1.1 (Martin, 2011). Reads were aligned to the reference genome (Bter_1.0, Refseq accession no. GCF_000214255.1 (Sadd *et al.*, 2015)) using STAR v2.5.2 (Dobin *et al.*, 2016) with standard parameters. Reads were counted per gene using HTseq v0.10.0 (Anders *et al.*, 2015). Differential expression analysis was then carried out after count normalisation via a negative binomial generalised linear model implemented by DEseq2 v1.20.0 (Love *et al.*, 2014) in R v.3.4.0 (http://www.R-project.org) with colony and reproductive status as independent variables. Genes were classed as differentially expressed when q<0.05 after correction for multiple testing using the Benjamini-Hochberg method (Benjamini and Hochberg, 1995).

Differential exon expression was determined using the DEXseq v1.26.0 R package (Anders *et al.*, 2012), briefly this package calculated the ratio of expression of a given exon in a given gene relative to other exons in the same gene for each sample. The relative exon expression per colony was then calculated taking into account dispersion between colonies. A GLM was then used to test for a difference in the relative proportion of expression of each exon between castes, accounting for sample differences and overall gene expression differences between castes, p-values were corrected for multiple testing using the Benjimini-Hochberg method (Benjamini and Hochberg, 1995) and exons were classed as differentially used between castes when q<0.1. The benefit of this method over general alternative splicing analysis is that specific differentially used exons can be identified between castes allowing the relationship with exonic methylation to be investigated.

### Differential methylation

BS-seq libraries were aligned to the reference genome (Bter_1.0, Refseq accession no. GCF_000214255.1, (Sadd *et al.*, 2015)) using Bismark v.0.16.1 (Krueger and Andrews, 2011) and bowtie2 v.2.2.6 (Langmead and Salzberg, 2012) with standard parameters. Bismark was also used to extract methylation counts and carry out deduplication. Annotation of the methylation counts with genomic features (from the *B. terrestis* annotation file, Refseq accession no. GCF_000214255.1) was carried out using custom R scripts implementing the sqldf v0.4.11 library (Grothendieck, 2017). Bisulfite conversion efficiency was calculated by aligning reads to the lambda reference genome (Refseq accession no. GCF_000840245.1) and calculating the single-site methylation level as in Schultz *et al.* (2012).

Prior to differential methylation analysis coverage outliers (above the 99.9th percentile) were removed along with bases covered by less than 10 reads. The methylation status of each CpG was then determined, using the ‘methylation status calling’ (MSC) procedure, as described in Cheng and Zhu (2014), briefly, this involves applying a mixed binomial model to each CpG which includes estimation of both the false discovery rate and the non-false discovery rate in order to make a binary methylation call per site. CpG sites were then filtered to remove any site that did not return as methylated in at least one sample. This functions to reduce the number of tests and hence decreases the stringency of the later FDR correction applied during differential methylation testing. This is a vital step for species with extremely low genome methylation where the majority of sites show zero methylation in all samples. A logistic regression model was then applied, via the R package methylKit v1.6.1 (Akalin *et al.*, 2012), to determine deferentially methylated sites, taking into account colony as a covariate due to high inter-colony variation, see supplementary 2.0. A minimum difference of at least 10% methylation and a q-value of <0.05 were required for a single site to be classed as differentially methylated. Genes containing at least one differentially methylated CpG and a minimum weighted methylated difference of 10% across the entire gene were classed as differentially methylated between reproductive castes.

We chose not to include a permutation test as part of the differential methylation analysis, as has been seen in previous research (Arsenault *et al.*, 2018; Libbrecht *et al.*, 2016), although it is included in our supplementary data. There is structure present in our data due to high methylation variation between colony replicates. When structure is present within data, permutation tests do not produce reliable outcomes, as discussed in Winkler *et al.* (2015). A higher number of replicates would allow label shuffling within confounding factors, maintaining the structure of the data, thus allowing a valid permutation, see supplementary 2.0.

### GO analysis

Gene ontology terms for *B. terrestris* were taken from a custom database made in Bebane *et al.* (2019). GO enrichment analysis was carried out using a hypergeometric test with Benjamini-Hochberg (Benjamini and Hochberg, 1995) multiple-testing correction. GO terms were defined as enriched when q<0.05. GO terms from differentially methylated genes were tested for enrichment against GO terms associated with all methylated genes. Genes were classed as methylated when the weighted methylation score per gene was greater than zero (Schultz *et al.*, 2012). Additionally the GO terms associated with hypermethylated genes in either sterile or reproductive workers were tested for enrichment against the GO terms associated with all differentially methylated genes between castes to determined if there are different functions for hypermethylated genes in either sterile or reproductive workers. GO terms for differentially expressed genes and genes containing different exon usage between castes were tested for enrichment against GO terms associated with all genes identified in the RNA-seq data. REVIGO (Supek *et al.*, 2011) was used to obtain the GO descriptions from the GO identification numbers.

### Comparative analyses

The hypergeometric test was applied to gene lists from the various analyses to determine if any overlap was statistically significant. Custom R scripts were used to investigate the relationship between gene expression and methylation. A reciprocal blast between the honeybee (Amel_4.5, Refseq accession no. GCA_000002195.1) and bumblebee genome (Bter_1.0, Refseq accession no. GCA_000214255.1) was carried out using blast+ v2.5.0 (Camacho *et al.*, 2009), where the fasta sequence for each gene of each species was blasted against a custom database containing the fasta sequence for every gene of the opposite species, allowing only one match per gene and a minimum e-value of 1 x10^-3^. The results were then filtered to ensure only matches that occurred in both directions and to only one gene were used. For example, multiple honeybee genes matched the same bumblebee gene, therefore all of these matches were discarded. This allowed us to construct a database of putative orthologous genes. A custom script was then used to check for overlap between the differentially methylated genes identified here and differentially methylated genes identified in Lyko *et al.* (2010) between honeybee reproductive castes.

## Results

### Genome-wide methylation differences between castes

Up to a maximum of 10bp were trimmed from the start of all reads due to base bias generated by the Illumina sequencing protocol (Krueger *et al.*, 2011). The mean mapping efficiency was 63.6% ± 1.4% (mean ± standard deviation) and the mean coverage was 17.7 ± 0.5 reads per base, the average number of uniquely mapped reads were 27,709,214 ± 753,203. The mean bisulfite conversion efficiency, calculated from the unmethylated lambda spike, was 99.55% ± 0.02%. After accounting for the conversion efficiency there were no methylated cytosines in a non-CpG context outside of this range. The mean single site methylation level (Schultz *et al.*, 2012) in a CpG context was determined as 0.22% ± 0.07, calculated from the number of methylated cytosines divided by the sum of methylated and unmethylated cytosines and accounting for bisulfite conversion efficiency.

A total of 3412 genes were classed as methylated, i.e. they had a weighted methylation level >0 in at least one sample. There was no significant difference in the overall weighted methylation level of the methylated genes between reproductive and sterile workers (Mann-Whitney U test: W = 5948300, p = 0.1172, Fig.3a). GO terms enriched in methylated genes compared to all genes annotated in the genome (q<0.05) include a large variety of biological processes (supplementary 1.0.5), specifically *post-transcriptional regulation of gene expression* (GO:0010608), *histone modification* (GO:0016570), *chromatin remodelling* (GO:0006338) are enriched as well as terms related to reproductive processes, e.g. *reproduction* (GO:0000003), *gamete generation* (GO:0007276), *oogenesis* (GO:0048477) and *oocyte differentiation* (GO:0009994).

**Figure 3:**
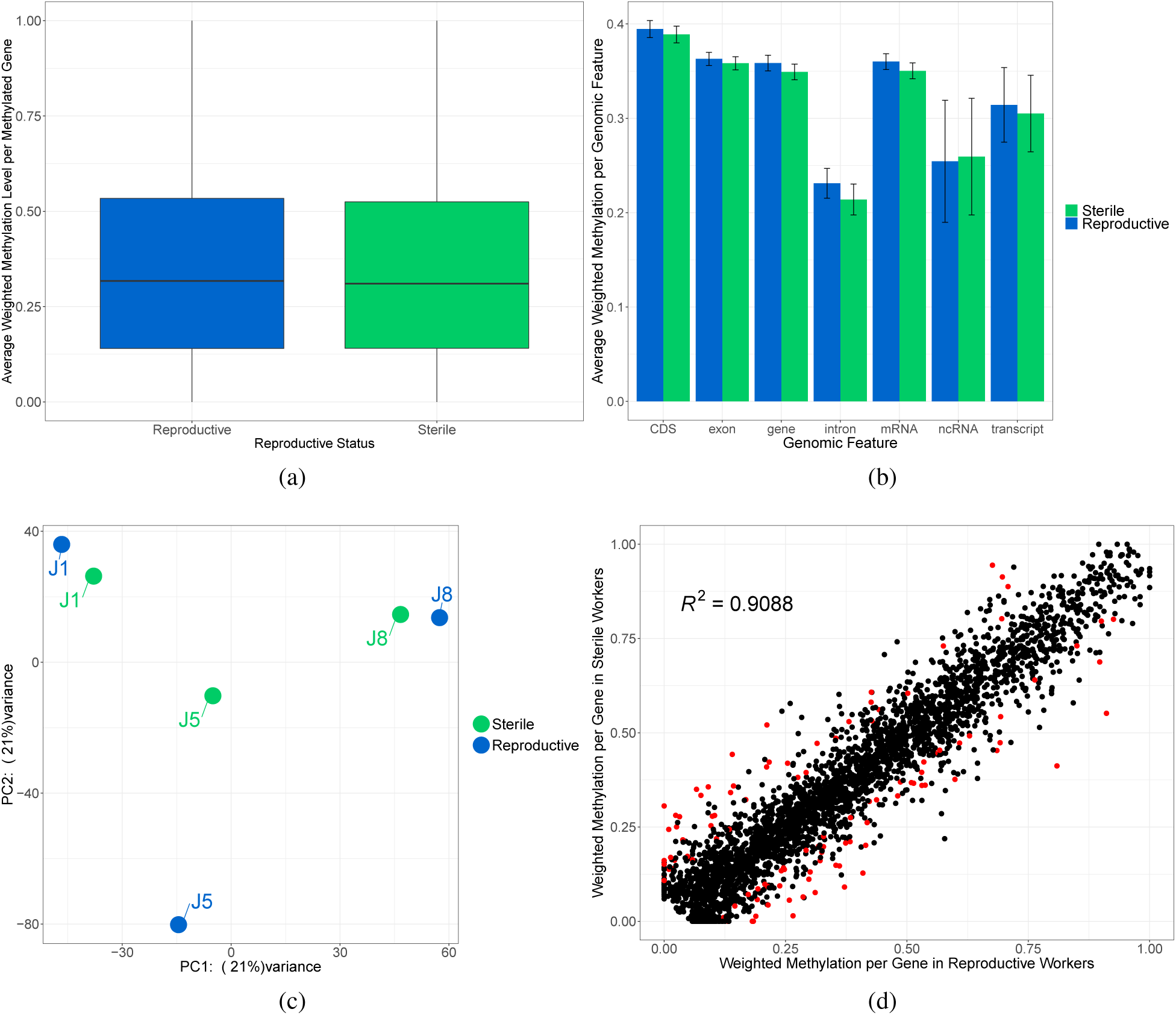
(a) Box plot of the mean weighted methylation level of methylated genes (n = 3412) across colonies for each caste. (b) The mean weighted methylation level across colonies for each genomic feature for both reproductive and sterile workers. Error bars are 95% confidence intervals of the mean. (c) PCA plot generated by methylKit showing samples cluster by colony using per site CpG methylation. (d) Scatter plot of the average weighted methylation level (across colonies) for reproductive workers against sterile workers. Each dot represents a gene and each red dot represents a differentially methylated gene (q<0.05 and a minimum gene weighted methylation difference of 10%).

There was no significant difference in the weighted methylation level of all genomic features between reproductive and sterile workers (Two-way ANOVA, interaction between genomic feature and reproductive status; F_1,6_ = 0.637, p = 0.701, Fig.3b), we also tested for differences in methylation levels of putative promotors but as promotor regions are currently unannotated for the current genomic annotation we feel these results are not reliable (supplementary 2.1, Fig.S3a). Irrespective of worker reproductive status we found methylation differences between genomic features (Kruskal-Wallis; chi-squared = 729.35, df = 7, p < 2.2 x 10^-16^). With methylation being significantly enriched in coding regions compared to introns and ncRNAs (supplementary 1.0.6).

We also found no difference in the weighted methylation level across the genome per linkage group between reproductive and sterile workers (Two-way ANOVA, interaction between linkage group and reproductive status; F_1,17_ = 0.034, p = 1.0, supplementary 2.1, Fig.S3b). Weighted methylation did vary significantly between linkage groups within the genome irrespective of reproductive status (Kruskal-Wallis chi-squared = 131.59, df = 17, p < 2.2 x 10^-16^, supplementary 1.0.6), however due to the number of unplaced scaffolds these results should be interpreted with care.

Finally using individual CpG methylation levels samples cluster by colony rather than reproductive caste (Fig.3c) indicating high inter-colony methylation variation.

### Gene level methylation differences between castes

A total of 4,681,131 CpG sites had a coverage >10 in all six sample data sets, of those 16,194 returned as methylated in at least one sample after running the MSC procedure. A total of 624 of these CpGs were identified as differentially methylated between reproductive castes, 613 of these were located in a total of 478 genes (supplementary 1.0.4). 11 differentially methylated CpGs were located outside of genes, nine of those were within 5000bp upstream or downstream of a gene with no apparent trend in the expression of near-by genes (supplementary 1.0.5). After stringent filtering of these genes to also require each gene to also show a minimum weighted methylation difference of 10% we found 111 differentially methylated genes between reproductive and sterile workers (supplementary 1.0.7, Fig.3d).

Of the 111 differentially methylated genes, there was no preference for genes to be hypermethylated in either reproductive or sterile workers (Chi-squared goodness of fit, X-squared = 2.027, df = 1, p = 0.1545), with 63 genes hypermethylated in reproductive workers and 48 genes hypermethylated in sterile workers.

GO terms enriched in differentially methylated genes compared to all methylated genes (q<0.05) contained a variety of biological processes (supplementary 1.0.8), amongst these processes were terms involved with reproduction, including; *meiotic cell cycle* (GO:0051321), *maintenance of oocyte nucleus location involved in oocyte dorsal/ventral axis specification* (GO:0042070), *negative regulation of transcription involved in meiotic cell cycle* (GO:0051038), *female meiosis chromosome segregation* (GO:0016321) and *female germline ring canal stabilisation* (GO:0008335). One of the genes associated with the above GO terms is *eggless* (LOC100647514: *histone-lysine N-methyltransferase eggless*) which shows hypermethylation in sterile workers. This gene contains a Methyl-CpG binding domain which has been associated with histone H3, lysine 9-specific methyltransferase which contributes to repression of transcription (Wakefield *et al.*, 1999).

There were no specific GO terms enriched for either the hypermethylated genes in sterile workers or the hypermethylated genes in reproductive workers compared to all differentially methylated genes as a background.

### Expression differences between castes

All reads had 13bp trimmed from the start due to base bias generated by the Illumina protocol (Krueger *et al.*, 2011). The mean number of uniquely mapped reads was 89.4% ± 0.8% (mean ± standard deviation). This equated to a mean of 10,115,366 ± 1,849,600 uniquely mapped reads. After running a differential expression analysis with DESeq2, the decision was made to remove one sample from all downstream analysis due to possible mislabelling of reproductive status (supplementary 2.2).

Samples cluster by reproductive status when the expression of all genes is assessed (Fig.4a). A total of 334 genes were identified as differentially expressed (q<0.05, Fig.4b). There was no difference in the number of up-regulated genes in either reproductive or sterile workers (Chi-squared goodness of fit: X-squared = 0.2994, df = 1, p-value = 0.5843), with 172 genes up-regulated in reproductive workers and 162 genes up-regulated in sterile workers.

**Figure 4:**
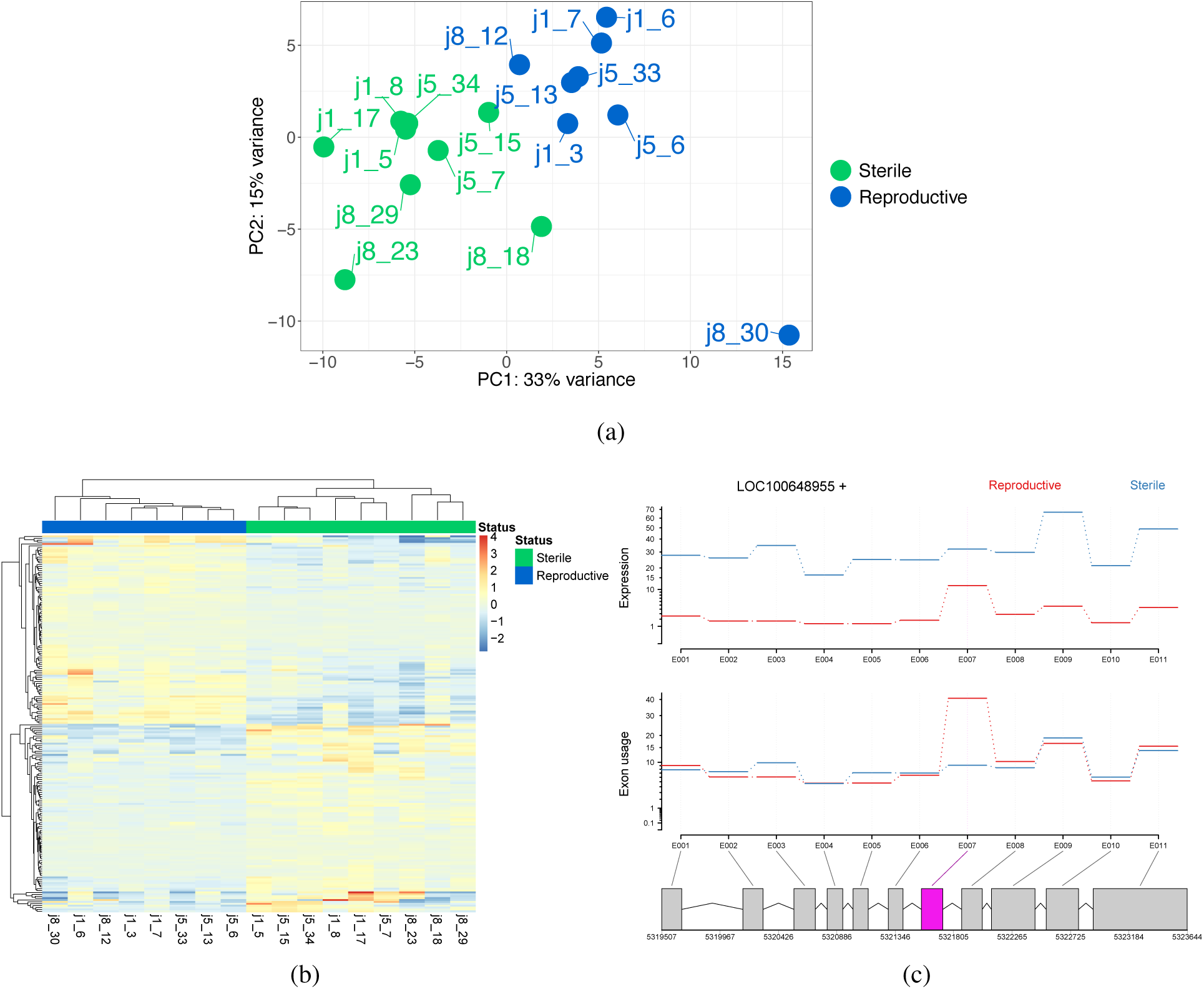
(a) PCA plot showing samples cluster by caste for gene expression, the first half of each label represents the colony name and the second half is the individual identification number. (b) Heatmap showing the 100 top differentially expressed genes between reproductive castes, samples cluster by reproductive status. Sample names are show at the bottom of the plot. (c) An example of a gene which shows differential exon expression in one exon between reproductive castes. The top section of the plot shows the general expression differences between castes, the second section shows the normalised counts per exon (given expression differences) and the third section highlights the differentially expressed exon in pink. Gene shown: *probable peroxisomal acyl-coenzyme A oxidase 1* (ID: LOC100648955).

One of the most up-regulated genes in reproductive workers was vitellogenin (gene ID: LOC100650436, log2 fold-change of 2.92, q = 4.85 x 10^-6^). Previous work has found up-regulation of this gene in reproductive *B. terrestris* workers is linked to aggressive behaviour rather than directly to ovary development (Amsalem *et al.*, 2014). Additionally two genes coding for serine-protease inhibitors were found to be up-regulated in reproductive workers, these proteins have been linked to reproduction in other insect species (Bao *et al.*, 2014).

Enriched GO terms associated with the differentially expressed genes compared to the background of all genes in the RNA-seq data (q<0.05) contained a variety of biological processes (supplementary 1.1.0), including *embryonic process involved in female pregnancy* (GO:0060136) and *positive regulation of oviposition* (GO:1901046). Additionally there were no specific GO terms enriched in up-regulated genes of reproductive workers compared to all differentially expressed genes as the background. However there were two GO terms enriched for up-regulated genes in sterile workers compared to differentially expressed genes as the background, these were; *cellular lipid metabolic process* (GO:0044255) and *isoprenoid biosynthetic process* (GO:0008299).

A total of 59 genes were identified as having differential exon usage, containing 83 differentially expressed exons between reproductive castes (q<0.1, supplementary 1.1.1, see example Fig.4c). There is no difference in the number of up-regulated exons in reproductive workers compared to sterile workers (Chi-squared goodness of fit: X-squared = 3.4819, df = 1, p-value = 0.06204), with reproductive workers having 33 up-regulated exons and sterile workers having 50 up-regulated exons. The enriched GO terms associated with genes containing differentially used exons compared to the background of all genes in the RNA-seq data (q<0.05) contained a variety of biological processes (supplementary 1.1.2.), however there were no GO terms with a clear connection to reproductive processes.

### Relationship of methylation and gene expression

On an individual gene basis, methylation and reproductive caste have no effect on expression level (Fig.5a and 5b, linear mixed effects model with colony as a random factor; methylation: df = 49172, t = −1.295, p = 0.195, reproductive status: df = 49172, t = −0.638, p = 0.524, interaction between methylation and reproductive status: df = 49172, t = 0.112, p = 0.911).

**Figure 5:**
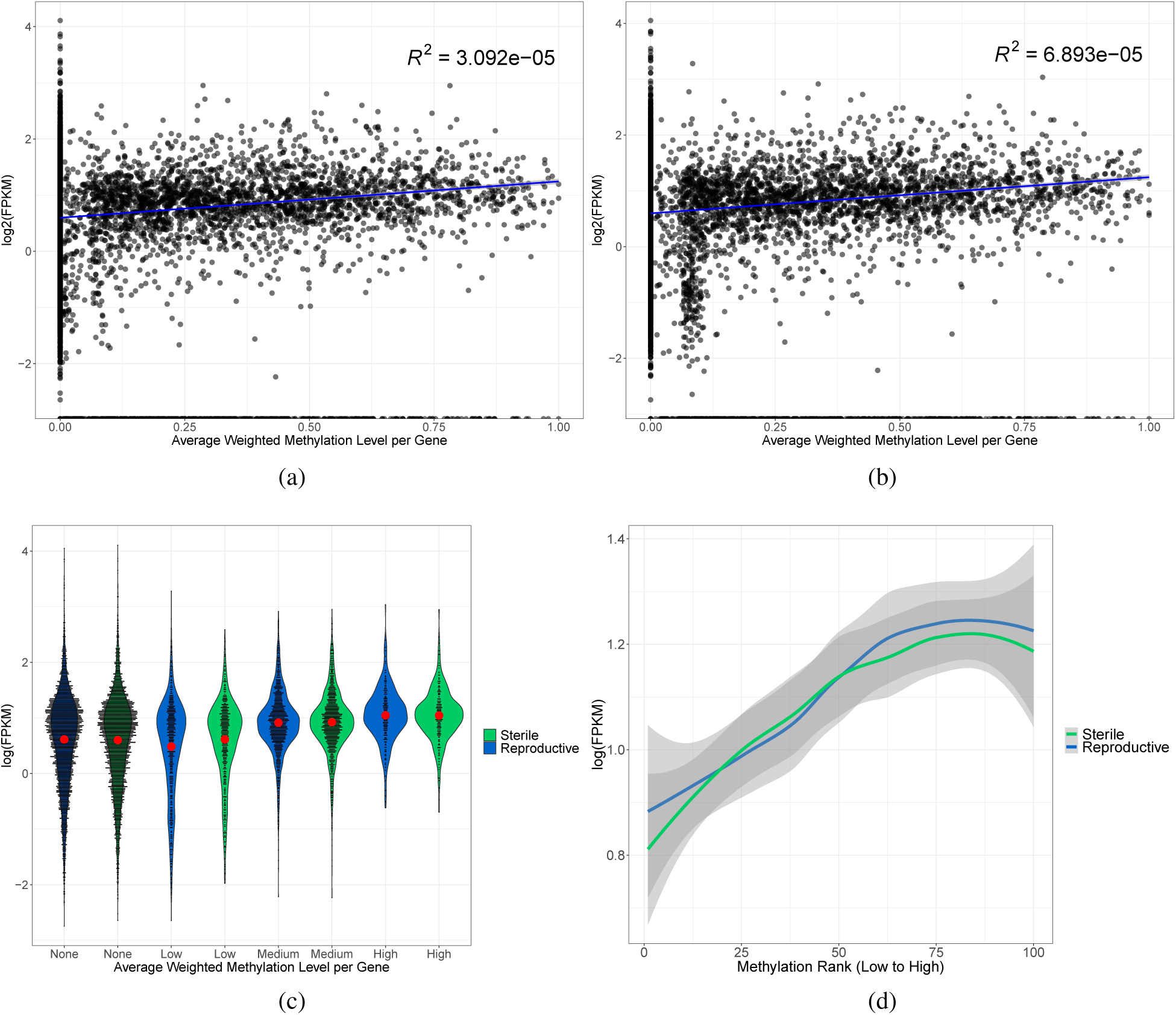
(a) and (b) The colony average weighted methylation level for every gene plotted against the log(FPKM) of that gene for sterile workers and reproductive workers respectively. Each black dot represents a single gene. The blue line is a fitted linear regression with the grey shaded area representing 95% confidence intervals. (c) Violin plots showing the distribution of the data via a mirrored density plot, meaning the widest part of the plots represent the most genes. Weighted methylation level per gene per caste, averaged across colonies, was binned into four categories, no methylation, low (>0 - 0.2), medium (0.2 - 0.7) and high (0.7 - 1), as in Liu *et al.* (2019). The red dot indicates the mean with 95% confidence intervals. Each black dot represents a single gene. (d) Binned methylated genes (n = 3412, 100 being the most highly methylated) based on the mean weighted methylation level across colonies per reproductive caste, plotted against the log(FPKM) expression level per gene. Data were smoothed using the LOESS method, grey areas are 95% confidence intervals.

However, gene groups with varying methylation levels show different levels of expression (Kruskal-Wallis; chi-squared = 131.59, df = 17, p < 2.2 x 10^-16^, Fig.5c). Specifically genes with no methylation show higher expression than genes classed as lowly methylated but lower expression than genes classed as medium/high in terms of methylation (Dunn’s test with Benjamini-Hochberg multiple-testing correction; no methylation vs low methylation: Z = −13.14 p = 4.09 x 10^-39^, no methylation vs medium methylation: Z = 4.5 p = 6.82 x 10^-6^, no methylation vs high methylation: Z = 7.32 p = 3.86 x 10^-13^, Fig.5c). Reproductive caste still has no effect on gene expression in relation to methylation status when genes are grouped (Two-way ANOVA, interaction between reproductive status and methylation level; F_1,3_ = 0.017, p = 0.99).

A linear mixed effects model was then applied to assess the relationship between gene expression, methylation and reproductive status for only methylated genes using colony as a random factor. There is a positive relationship between gene expression and methylation in methylated genes with reproductive status having no effect (Fig.5d, methylation: df = 17390, t = 6.154, p = 7.72 x 10^-10^, reproductive status: df = 17390, t = −0.328, p = 0.743, interaction between methylation and reproductive status: df = 17390, t = −0.200, p = 0.842).

### Relationship of methylation and differential gene expression

Weighted methylation differences between differentially expressed genes and non-differentially expressed genes were assessed along with weighted methylation differences between genes containing differentially expressed exons and genes without differentially expressed exons (Fig.6a and Fig.6b). Differentially expressed genes and genes containing differentially expressed exons between castes show lower methylation than non-differentially expressed genes or genes containing no differentially expressed exons, with reproductive status and the interaction of reproductive status with gene expression type having no effect (Table 1, Fig.6a and 6b). When weighted methylation is assessed per exon, differentially expressed exons have lower weighted methylation than non-differentially expressed exons (Fig.6c), with reproductive status and the interaction of reproductive status and exon expression having no effect (Table 1).

**Figure 6:**
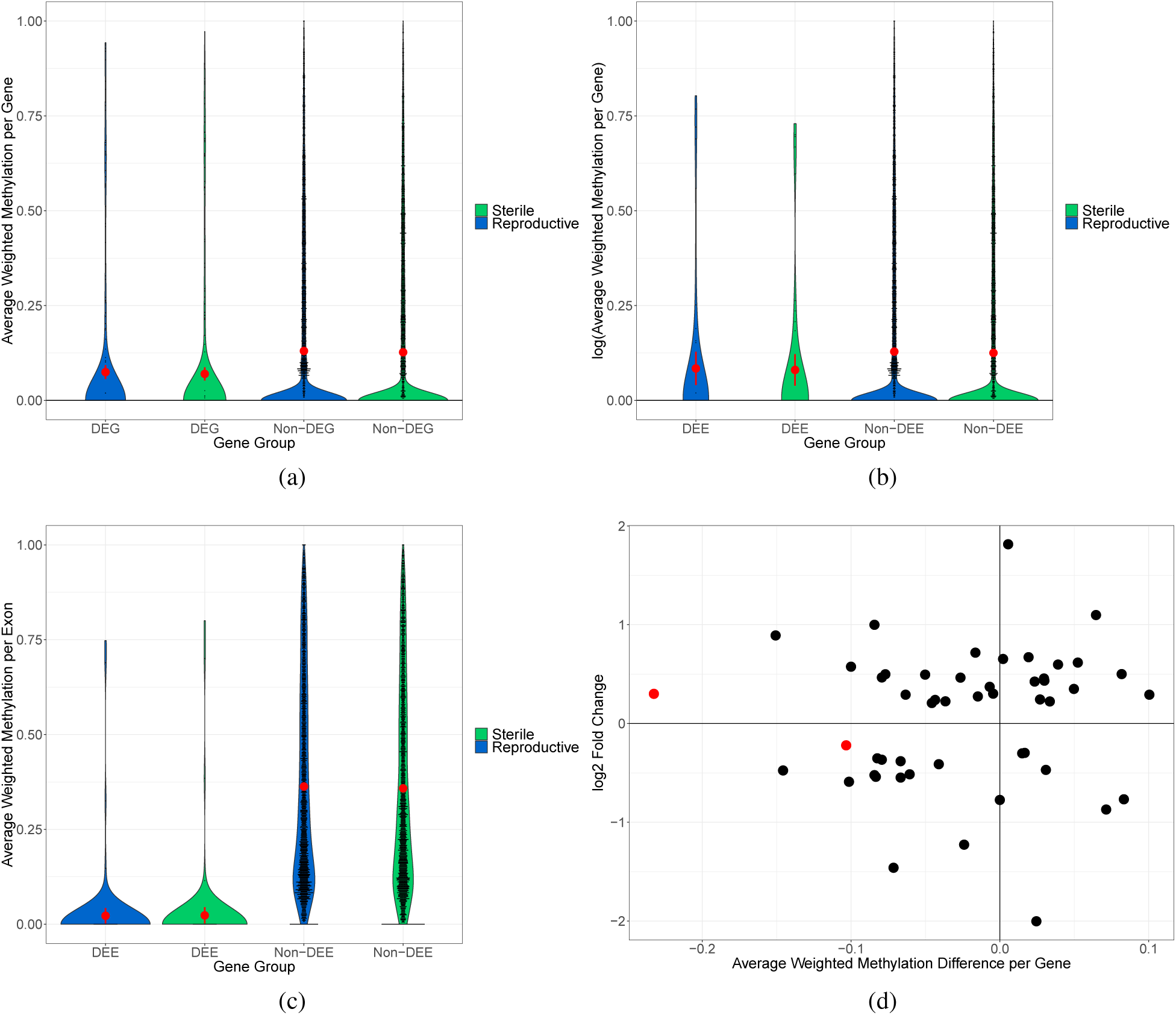
(a) Violin plots showing the distribution of the data via a mirrored density plot, meaning the widest part of the plots represent the most genes. The red dots represent the mean of each gene set along with error bars representing 95% confidence intervals of the mean. Each black dot is an individual gene. The mean weighted methylation per gene across colonies per caste is plotted for either differentially expressed genes (DEG) or non-differentially expressed genes (non-DEG). (b) Violin plots of the mean weighted methylation per gene across colonies per caste is plotted for either genes containing differentially expressed exons (DEE) or genes with no differentially expressed exons (non-DEE). (c) Violin plots of the mean weighted methylation per exon across colonies per caste for differentially expressed exons (DEE) and non-differentially expressed exons (non-DEE). The red dots represent the mean of each gene set along with error bars representing 95% confidence intervals of the mean. Each black dot is an individual exon. (d) Scatter plot of the difference in the mean weighted methylation level across colonies between castes plotted against the log2 fold change in expression of differentially expressed genes between castes. Each dot represents a gene, only genes which have a methylation difference >0 are shown. The red dots indicate the gene is also differentially methylated.

**Table 1:**
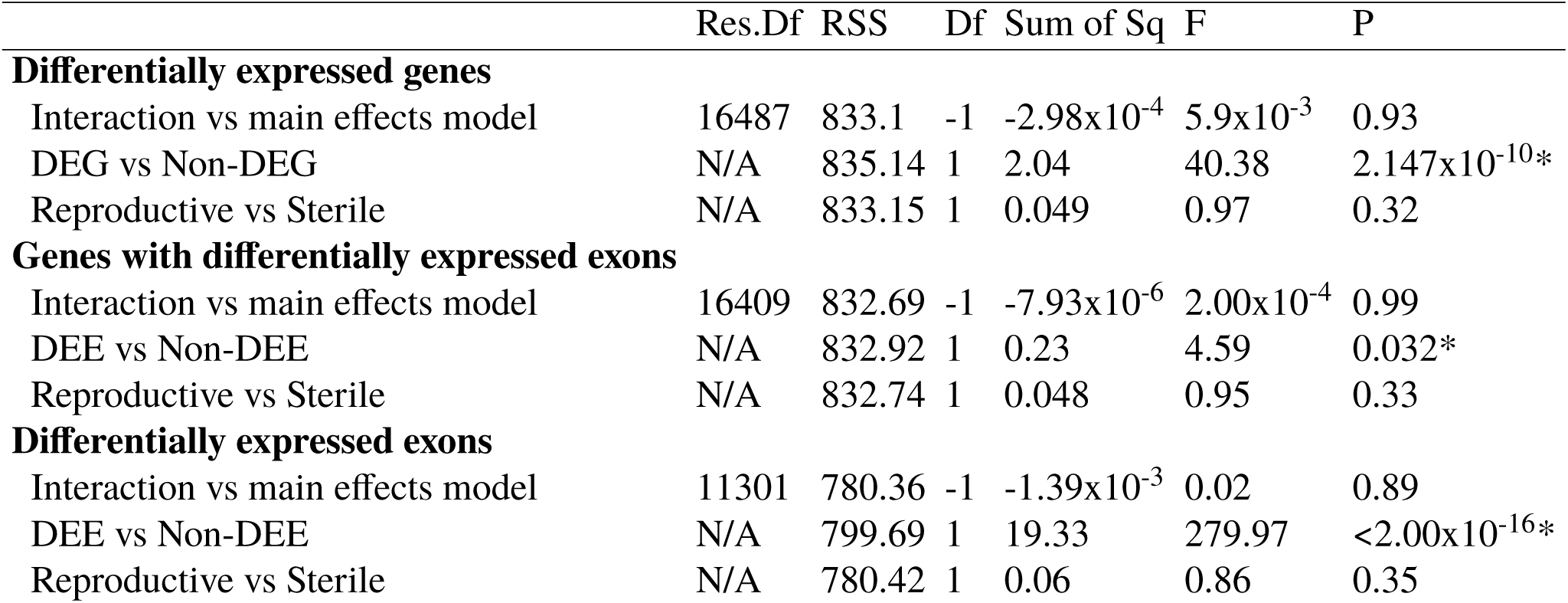
Summary statistics of the linear models used to check for differences in weighted methylation level between gene sets, taking into account reproductive status. The interaction vs main effects models were tested using the *anova* function in R to assess the interaction effect between gene set and reproductive status. * indicates a significant p-value <0.05.

|  | Res.Df | RSS | Df | Sum of Sq | F | P |
| --- | --- | --- | --- | --- | --- | --- |
| **Differentially expressed genes** |  |  |  |  |  |  |
| Interaction vs main effects model | 16487 | 833.1 | -1 | -2.98x10<sup>-4</sup> | 5.9x10<sup>-3</sup> | 0.93 |
| DEG vs Non-DEG | N/A | 835.14 | 1 | 2.04 | 40.38 | 2.147x10<sup>-10</sup>* |
| Reproductive vs Sterile | N/A | 833.15 | 1 | 0.049 | 0.97 | 0.32 |
| **Genes with differentially expressed exons** |  |  |  |  |  |  |
| Interaction vs main effects model | 16409 | 832.69 | -1 | -7.93x10<sup>-6</sup> | 2.00x10<sup>-4</sup> | 0.99 |
| DEE vs Non-DEE | N/A | 832.92 | 1 | 0.23 | 4.59 | 0.032* |
| Reproductive vs Sterile | N/A | 832.74 | 1 | 0.048 | 0.95 | 0.33 |
| **Differentially expressed exons** |  |  |  |  |  |  |
| Interaction vs main effects model | 11301 | 780.36 | -1 | -1.39x10<sup>-3</sup> | 0.02 | 0.89 |
| DEE vs Non-DEE | N/A | 799.69 | 1 | 19.33 | 279.97 | <2.00x10<sup>-16</sup>* |
| Reproductive vs Sterile | N/A | 780.42 | 1 | 0.06 | 0.86 | 0.35 |

Of the 334 differentially expressed genes, 50 also showed some level of weighted methylation difference between reproductive and sterile workers (weighted methylation difference >0), Fig.6d. However, there is no relationship between the level of differential methylation and the level of differential expression for these 50 genes (linear model: F_1, 58_ = 0.2717, p = 0.6046).

Gene lists were checked for potential overlap from all analyses. There was no significant overlap between differentially methylated genes and differentially expressed genes (two genes, hypergeometric test; p = 0.658, Fig.7a). There was also no significant overlap between differentially methylated genes and genes containing differentially expressed exons (one gene, hypergeometric test; p = 0.12, Fig.7a).

**Figure 7:**
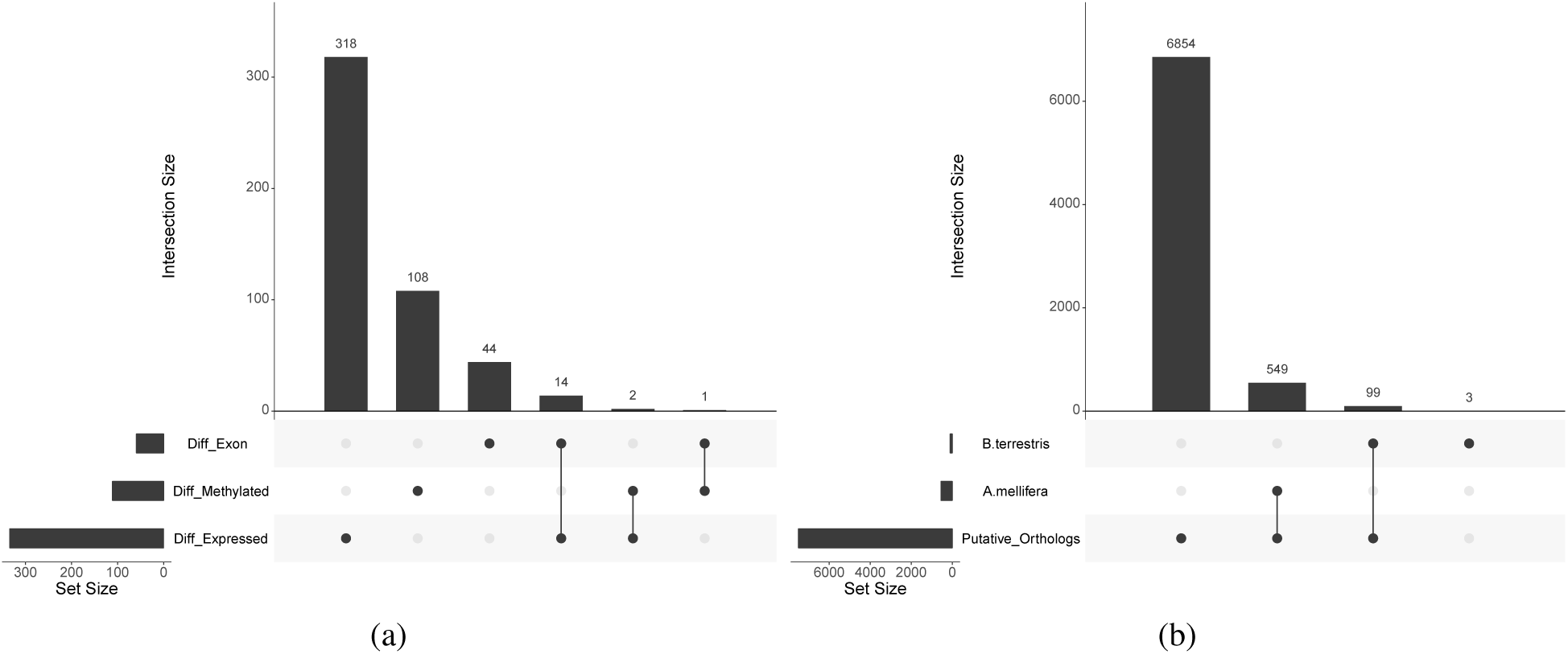
(a) UpSet plot showing the number of common genes between analyses. The set size indicates the number of genes in each category; differentially expressed, differentially methylated or genes containing a differentially expressed exon. The intersection size indicates the number of genes either unique to each set or the number common between sets. A single dot in the lower panel indicates the number of genes unique to the corresponding set and joining dots indicate the number of genes in common between the corresponding sets. (b) UpSet plot showing the number of putative orthologs between *A. mellifera* and *B. terrestris* along with the number of differentially methylated genes identified in Lyko *et al.* (2010) and in this study which are present in the putative ortholog database.

There was a significant overlap of genes found to be differentially expressed with those containing differentially expressed exons, 14 total (hypergeometric test; p = 1.44 10^-10^, Fig.7a). All lists of overlapping genes can be found in supplementary 1.1.3.

### Honeybee orthologous differentially methylated genes

Custom honeybee and bumblebee putative-ortholog databases were created from 15,314 and 10,339 annotated genes respectively (Amel_4.5 GCA_000002195.1, Bter_1.0 GCA_000214255.1). 9,244 honeybee genes matched at least one bumblebee gene and 7,985 bumblebee genes matched at least one honeybee gene with an e-value of <1 x 10^-3^. A total of 7,345 genes made the same match in both blast searches. Of these genes 392 matched more than one gene in one or both blasts and were therefore removed. This left a final putative ortholog list of 6,953 genes. 99 of the 111 differentially methylated genes identified here were present in the final putative ortholog list however none of them matched the 549 genes identified as differentially methylated between honeybee reproductive castes by Lyko *et al.* (2010), Fig.7b.

## Discussion

We have used whole genome bisulfite sequencing and gene expression libraries from the same individual *B. terrestris* workers to investigate the role of methylation in caste determination. We found both reproductive and sterile workers show similar methylation patterns to other social insects. Methylation also has a similar relationship with gene expression compared to most other social insects currently studied, with more highly methylated genes showing higher levels of expression and lower levels of methylation being associated with differentially expressed genes. We found no methylation differences on a genomic scale between castes, however 111 genes were differentially methylated, these were involved in a variety of functions including reproductive related processes. We found no relationship between genes that are differentially expressed, or contain differential exon usage between castes with those that show differential methylation between castes. Finally we also found no common putative orthologous genes differentially methylated between *B. terrestris* and *A. mellifera* reproductive castes.

This is the first data set to accurately quantify methylation at base-pair resolution for *B. terrestris*. It confirms low methylation levels throughout the genome as predicted by Sadd *et al.* (2015). These low levels along with the enrichment for CpG methylation in coding regions is also seen in many social insect species, including *A. mellifera* (Lyko *et al.*, 2010) and multiple ant species (Bonasio *et al.*, 2012; Libbrecht *et al.*, 2016). However this trend is not completely conserved among all social insects, for example the primitively social wasp species *Polistes dominula* shows 6% CpG methylation (Weiner *et al.*, 2013), and the highly social termite, *Zootermopsis nevadensis*, has exceptionally high methylation levels compared to the majority of insects (Bewick *et al.*, 2017), with 12% CpG methylation, and methylation being just as common in introns as exons (Glastad *et al.*, 2016).

Higher levels of CpG methylation are associated with higher levels of gene expression in both *B. terrestris* reproductive castes. This is also the case in other social insects, with Figures 5a and 5d showing almost identical trends to those found in Bonasio *et al.* (2012), Patalano *et al.* (2015) and Libbrecht *et al.* (2016). Additionally other social insect species, show higher methylation in non-differentially expressed genes as we found here, examples include: *Dinoponera quadriceps* (Patalano *et al.*, 2015), *Polistes canadensis* (Patalano *et al.*, 2015), *Zootermopsis nevadensis* (Glastad *et al.*, 2016) and *Cerapachys biroi* (Libbrecht *et al.*, 2016). Higher levels of methylation in more highly expressed genes and in non-differentially expressed genes is thought to indicate a role for methylation in housekeeping genes in social insects (Foret *et al.*, 2009; Lyko *et al.*, 2010; Bonasio *et al.*, 2012; Wang *et al.*, 2013). High levels of gene body methylation is also found in highly expressed genes in plants, whilst currently the function is unknown, Zilberman (2017) predicts it functions to stabilise expression by reducing histone variants and Bewick and Schmitz (2017) predict it is a byproduct of transposable element silencing.

As well as a similar methylation profile to other social insects, we also predicted that if methylation plays a role in reproductive caste determination we would find differentially methylated genes between castes with reproductive related functions. We found 111 differentially methylated genes which include enriched GO terms for various reproductive related processes, this suggests methylation has some association with the switch between sterility and reproduction in *B. terrestris*. This supports previous work which exposed *B. terrestris* to a chemical which decreases general methylation levels and found workers were more likely to become reproductive (Amarasinghe *et al.*, 2014), however we did not find a difference in the genome-wide methylation levels of sterile and reproductive workers. It is also worth noting a worker classed as reproductive appeared to show a sterile transcriptional profile and this was included in the pool for the reproductive sample for colony J8. This will have ‘diluted’ the strength of the methylation profile for this particular sample. It is therefore likely our data contains false negatives, meaning there may be differentially methylated genes between reproductive castes which do not appear in our data set.

Whilst we found differentially methylated genes between castes we found no evidence for methylation directly affecting gene expression between reproductive and sterile workers. Only a small non-significant number of genes are both differentially methylated and differentially expressed between castes and there is no relationship between the degree of differential methylation and differential expression on a gene level. Previous research using the milkweed bug (*Oncopeltus fasciatus*) found knocking down *Dnmt1*, the gene responsible for DNA methylation maintenance, in ovary tissue with RNAi had no effect on gene expression, however these individuals could no longer reproduce (Bewick *et al.*, 2019). This suggests methylation may play an alternative role, rather than direct regulation of gene expression, in reproduction of some insects. Bewick *et al.* (2019) suggest this role may be the regulation of genome stability and/or the regulation of vital cellular processes. The variety of GO terms involved in biological processes we obtained for the differentially methylated genes between castes supports this idea.

Additionally, we observed high inter-colony variation in methylation, however we did not have sufficient replicates to test for exact differences between colonies. High inter-colony variation could suggest methylation may also play a role in the adaptive abilities of *B. terrestris*. Stable environmentally induced ‘epialleles’ have been proposed to act as an additional layer of information for which selection can act upon (Flores *et al.*, 2013). However we currently do not know if *B. terrestris* methylation shows trans-generational inheritance or whether a large proportion is wiped during development, as in mammals (Messerschmidt *et al.*, 2014).

It has also been suggested methylation may regulate alternative splicing, rather than expression in some insects (Glastad *et al.*, 2011). We found no evidence here for the role of exon methylation in exonic expression differences between castes. Previous research using honeybees however did find an association between methylation and caste specific alternative splicing. Methylation differences between queens and workers in *A. mellifera* have been associated with caste specific splicing events (Lyko *et al.*, 2010). Additionally a knock-down of *Dnmt3* by RNA interference was found to affect alternative splicing patterns in *A. mellifera*, with decreased methylation levels being directly related to exon skipping and intron retention (Li-Byarlay *et al.*, 2013). However another social insect species, the primitively social wasp, *Polistes dominula*, has also shown no direct association between methylation and alternative splicing (Standage *et al.*, 2016), importantly this species shows extremely low genome methylation and appears to lack the *Dnmt3* gene responsible for *de novo* methylation, suggesting that the link between methylation and alternative splicing in social insects is variable.

Exon methylation has been shown to play a role in histone modifications and nucleosome stability in mammals (Singer *et al.*, 2015; Jones, 2012). These modifications have the ability to affect alternative splicing patterns through RNA polymerase accessibility, meaning whilst changes in DNA methylation may not be observed as directly related to alternative splicing, it’s possible these changes have a downstream effect leading to transcriptional changes (Hunt *et al.*, 2013). The analysis of the relationship between methylation and alternative splicing done here could be elaborated on further to include non-caste specific splicing sites and to also potentially identify the role of exon methylation in other epigenetic processes, which may themselves, lead to alternative splicing.

It is also worth noting other epigenetic mechanisms may play a role in caste determination, for example microRNAs have been associated with caste switching in *A. mellifera* (Ashby *et al.*, 2016). Additionally Simola *et al.* (2016) found histone acetylation differences between worker castes of *Camponotus floridanus*. They inhibited histone acetylation and found this caused the major worker caste to behave more like a minor worker. This same species has also been shown to have caste specific methylation profiles (Bonasio *et al.*, 2012). These examples indicate it is likely an interplay between multiple mechanisms that ultimately cause social insect caste differentiation, again supported by the fact we find no association between methylation and caste-specific alternative splicing.

Our final prediction was that differentially methylated genes between worker *B. terrestris* castes would be similar to those found to be differentially methylated between *A. mellifera* reproductive castes if methylation was involved in caste determination in Hymenoptera; we did not find any putative orthologues in common. This supports the idea in Bewick *et al.* (2019) that methylation may not directly influence caste determination. However the differentially methylated gene list obtained for *A. mellifera* used queen samples to represent the reproductive caste (Lyko *et al.*, 2010), whereas here reproductive worker samples were used, this could also explain the lack of agreement.

Considerably more experimental research is needed to better define the relationship between epigenetic processes and caste determination in social insects. Future work should focus on the consequences of experimental methylation removal or addition (Pegoraro *et al.*, 2017), as well as exploring additional epigenetic mechanisms to attempt to identify a full pathway leading to reproductive caste differences. For example, CRISPR has recently been used to knockout two sex-determining genes in *A. mellifera* causing individuals to change gender (Mcafee *et al.*, 2019). This technology has also been adapted to be able to change the methylation state of a given loci (Vojta *et al.*, 2016), allowing the possibility of exploring the function of methylation in specific genes.

Overall, we have found the *B. terrestris* methylome appears similar to some other social insects, in terms of overall levels and the relationship with gene expression. We found no methylation differences genome-wide between reproductive castes however we did find differentially methylated genes between reproductive castes, with GO terms enriched in many biological processes including reproduction. These results combined with previous research, (Amarasinghe *et al.*, 2014), indicate an association between methylation and reproductive caste differences in *B. terrestris*. However it is clear, owing to the lack of consistency between differentially methylated genes and differentially expressed genes, methylation is not directly responsible for the associated changes in gene expression leading to the different reproductive phenotypes in *B. terrestris*. Additionally the lack of similarity between differentially methylated genes between castes in *B. terrestris* and between castes in *A. mellifera* suggest methylation may not directly contribute to caste determination in some Hymenoptera. Future work should focus on the experimental manipulation of epigenetic processes, such as methylation, in social insects in order to clarify functional roles within and across species.

## Acknowledgements

Thank you to Dr. Ben Hunt and Boris Berkhout for important advice and discussion regarding data analysis. Thank you to Dr. Ezio Rosato for providing valuable discussion. This research used the ALICE2 High Performance Computing Facility at the University of Leicester. H.M. was supported by a NERC CENTA DTP studentship. Z.N.L. was supported by a BBSRC MIBTP DTP studentship. E.B.M. was funded by NERC grant NE/N010019/1. The authors delcare no conflict of interests.

## Author contributions

E.B.M. conceived the study. H.M. and Z.N.L. conducted the experiment. H. M. analysed the data and wrote the initial manuscript. All authors contributed to and reviewed the final manuscript.

## Data Accessibility

Data has been deposited in GenBank under NCBI BioProject: PRJNA533306. All code will be also be made available at: DOI:10.5281/zenodo.2394171.

## References

Akalin, A., Kormaksson, M., Li, S., Garrett-bakelman, F. E., Figueroa, M. E., Melnick, A., and Mason, C. E. 2012. methylKit: a comprehensive R package for the analysis of genome-wide DNA methylation profiles. Genome Biology, 13(R87).

Alaux, C., Jaisson, P., and Hefetz, A. 2006. Regulation of worker reproduction in bumblebees (*Bombus terrestris*): workers eavesdrop on a queen signal. Behavioral Ecology and Sociobiology, 60(3): 439–446.

Amarasinghe, H. E., Clayton, C. I., and Mallon, E. B. 2014. Methylation and worker reproduction in the bumble-bee (*Bombus terrestris*). Proceedings of the Royal Society B: Biological Sciences, 281(20132502).

Amsalem, E., Malka, O., Grozinger, C., and Hefetz, A. 2014. Exploring the role of juvenile hormone and vitellogenin in reproduction and social behavior in bumble bees. BMC Evolutionary Biology, 14(1): 45.

Anders, S., Reyes, A., and Huber, W. 2012. Detecting differential usage of exons from RNA-seq data. Genome Research, 22(10): 2008–2017.

Anders, S., Pyl, P. T., and Huber, W. 2015. HTSeq-A Python framework to work with high-throughput sequencing data. Bioinformatics, 31(2): 166–169.

Arsenault, S. V., Hunt, B. G., and Rehan, S. M. 2018. The effect of maternal care on gene expression and DNA methylation in a subsocial bee. Nature Communications, 9(3468).

Ashby, R., Forêt, S., Searle, I., and Maleszka, R. 2016. MicroRNAs in Honey Bee Caste Determination. Scientific Reports, 6(18794).

Bao, Y. Y., Qin, X., Yu, B., Chen, L. B., Wang, Z. C., and Zhang, C. X. 2014. Genomic insights into the serine protease gene family and expression profile analysis in the planthopper, Nilaparvata lugens. BMC Genomics, 15(1).

Bebane, P., Hunt, B. J., Pegoraro, M., Jones, A., Marshall, H., Rosato, E., and Mallon, E. 2019. The effects of the neonicotinoid imidacloprid on gene expression and DNA methylation in the buff-tailed bumblebee *Bombus terrestris*. bioRxiv, (http://dx.doi.org/10.1101/590091).

Benjamini, Y. and Hochberg, Y. 1995. Controlling the false discovery rate: a practical and powerful approach to multiple testing. Journal of the Royal Statistical Society, 57(1): 289–300.

Bewick, A. J. and Schmitz, R. J. 2017. Gene body DNA methylation in plants. Current Opinion in Plant Biology, 36: 103–110.

Bewick, A. J., Vogel, K. J., Moore, A. J., and Schmitz, R. J. 2017. Evolution of DNA methylation across insects. Molecular Biology and Evolution, 34(3): 654–665.

Bewick, A. J., Sanchez, Z., McKinney, E. C., Moore, A. J., Moore, P. J., and Schmitz, R. J. 2019. Dnmt1 is essential for egg production and embryo viability in the large milkweed bug, *Oncopeltus fasciatus*. Epigenetics and Chromatin, 12(1): 1–14.

Bloch, G. 1999. Regulation of queen-worker conflict in bumble-bee (*Bombus terrestris*) colonies. Proc. R. Soc. Lond. B, (266): 2465–2469.

Bonasio, R., Li, Q., Lian, J., Mutti, N. S., Jin, L., Zhao, H., Zhang, P., Wen, P., Xiang, H., Ding, Y., Jin, Z., Shen, S. S., Wang, Z., Wang, W., Wang, J., Berger, S. L., Liebig, J. J., Zhang, G., and Reinberg, D. 2012. Genome-wide and caste-specific DNA methylomes of the ants *Camponotus floridanus* and *Harpegnathos saltator*. Current Biology, 22(19): 1755–1764.

Camacho, C., Coulouris, G., Avagyan, V., Ma, N., Papadopoulos, J., Bealer, K., and Madden, T. L. 2009. BLAST+: Architecture and applications. BMC Bioinformatics, 10: 1–9.

Cheng, L. and Zhu, Y. 2014. A classification approach for DNA methylation profiling with bisulfite next-generation sequencing data. Bioinformatics, 30(2): 172–179.

Chevin, L.-M., Lande, R., and Mace, G. M. 2010. Adaptation, Plasticity, and Extinction in a Changing Environment: Towards a Predictive Theory. PLoS Biology, 8(4): e1000357.

Collins, D. H., Mohorianu, I., Beckers, M., Moulton, V., Dalmay, T., and Bourke, A. F. 2017. MicroRNAs Associated with Caste Determination and Differentiation in a Primitively Eusocial Insect. Scientific Reports, 7(45674).

Dobin, A., Gingeras, T. R., and Spring, C. 2016. Mapping RNA-seq Reads with STAR Alexander. Current Protocols in Bioinformatics, (51): 1–11.

Duchateau, M. J. and Velthuis, H. H. W. 1988. Development and reproductive strategies in *Bombus terrestris* colonies. Behaviour, 107(3): 186–207.

Fang, F., Hodges, E., Molaro, a., Dean, M., Hannon, G. J., and Smith, a. D. 2012. Genomic landscape of human allele-specific DNA methylation. Proceedings of the National Academy of Sciences, 109(19): 7332–7337.

Feng, S., Cokus, S. J., Zhang, X., Chen, P.-Y., Bostick, M., Goll, M. G., Hetzel, J., Jain, J., Strauss, S. H., Halpern, M. E., Ukomadu, C., Sadler, K. C., Pradhan, S., Pellegrini, M., and Jacobsen, S. E. 2010. Conservation and divergence of methylation patterning in plants and animals. Proceedings of the National Academy of Sciences, 107(19): 8689–8694.

Flores, K. B., Wolschin, F., and Amdam, G. V. 2013. The role of methylation of DNA in environmental adaptation. Integrative and Comparative Biology, 53(2): 359–372.

Foret, S., Kucharski, R., Pittelkow, Y., Lockett, G. A., and Maleszka, R. 2009. Epigenetic regulation of the honey bee transcriptome: unravelling the nature of methylated genes. BMC Genomics, 10(1): 472.

Foster, R. L., Brunskill, A., Verdirame, D., and O’Donnell, S. 2004. Reproductive physiology, dominance interactions, and division of labour among bumble bee workers. Physiological Entomology, 29(4): 327–334.

Geva, S., Hartfelder, K., and Bloch, G. 2005. Reproductive division of labor, dominance, and ecdysteroid levels in hemolymph and ovary of the bumble bee *Bombus terrestris*. Journal of Insect Physiology, 51(7): 811–823.

Glastad, K. M., Hunt, B. G., Yi, S. V., and Goodisman, M. A. 2011. DNA methylation in insects: On the brink of the epigenomic era. Insect Molecular Biology, 20(5): 553–565.

Glastad, K. M., Chau, L. M., and Goodisman, M. A. 2015. Epigenetics in Social Insects. In Physiology, Behavior, Genomics of Social Insects, volume 48, pages 227–269. Elsevier Ltd., 1 edition.

Glastad, K. M., Gokhale, K., Liebig, J., and Goodisman, M. A. D. 2016. The caste- and sex-specific DNA methylome of the termite *Zootermopsis nevadensis*. Scientific Reports, 6(37110).

Grothendieck, G. 2017. sqldf: Manipulate R Data Frames Using SQL. R package: https://cran.r-project.org/package=sqldf;

Herb, B. R., Wolschin, F., Hansen, K. D., Aryee, M. J., Langmead, B., Irizarry, R., Amdam, G. V., and Feinberg, A. P. 2012. Reversible switching between epigenetic states in honeybee behavioral subcastes. Nature Neuroscience, 15(10): 1371–1373.

Hunt, B. G., Glastad, K. M., Yi, S. V., and Goodisman, M. A. D. 2013. The function of intragenic DNA methylation: Insights from insect epigenomes. Integrative and Comparative Biology, 53(2): 319–328.

Jones, P. A. 2012. Functions of DNA methylation: Islands, start sites, gene bodies and beyond. Nature Reviews Genetics, 13(7): 484–492.

Krueger, F. and Andrews, S. R. 2011. Bismark: A flexible aligner and methylation caller for Bisulfite-Seq applications. Bioinformatics, 27(11): 1571–1572.

Krueger, F., Andrews, S. R., and Osborne, C. S. 2011. Large scale loss of data in low-diversity illumina sequencing libraries can be recovered by deferred cluster calling. PLoS ONE, 6(1): 4–10.

Kucharski, R., Maleszka, J., Foret, S., and Maleszka, R. 2008. Nutritional control of reproductive status in honeybees via DNA methylation. Science, 319(5871): 1827–1830.

Langmead, B. and Salzberg, S. L. 2012. Fast gapped-read alignment with Bowtie 2. Nat Methods, 9(4): 357–359.

Li, B., Hou, L., Zhu, D., Xu, X., An, S., and Wang, X. 2018. Identification and caste-dependent expression patterns of DNA methylation associated genes in *Bombus terrestris*. Scientific Reports, 8(2332).

Li-Byarlay, H., Li, Y., Stroud, H., Feng, S., Newman, T. C., Kaneda, M., Hou, K. K., Worley, K. C., Elsik, C. G., Wickline, S. A., Jacobsen, S. E., Ma, J., and Robinson, G. E. 2013. RNA interference knockdown of DNA methyl-transferase 3 affects gene alternative splicing in the honey bee. Proceedings of the National Academy of Sciences, 110(31): 12750–12755.

Libbrecht, R., Oxley, P. R., Keller, L., and Kronauer, D. J. C. 2016. Robust DNA Methylation in the Clonal Raider Ant Brain. Current Biology, 26: 1–5.

Liu, S., Aageaard, A., Bechsgaard, J., and Bilde, T. 2019. DNA Methylation Patterns in the Social Spider, *Stegodyphus dumicola*. Genes, 10(2): 137.

Lonsdale, Z., Lee, K., Kiriakidu, M., Amarasinghe, H., Nathanael, D., O’Connor, C. J., and Mallon, E. B. 2017. Allele specific expression and methylation in the bumblebee, *Bombus terrestris*. PeerJ, 5: e3798.

Love, M. I., Huber, W., and Anders, S. 2014. Moderated estimation of fold change and dispersion for RNA-seq data with DESeq2. Genome biology, 15(12): 550.

Lyko, F., Foret, S., Kucharski, R., Wolf, S., Falckenhayn, C., and Maleszka, R. 2010. The honey bee epigenomes: Differential methylation of brain DNA in queens and workers. PLoS Biology, 8(11).

Martin, M. 2011. Cutadapt removes adapter sequences from high-throughput sequencing reads. EMBnet.journal, 17(1): 10.

Matsuura, K., Mizumoto, N., Kobayashi, K., Nozaki, T., Fujita, T., Yashiro, T., Fuchikawa, T., Mitaka, Y., and Vargo, E. L. 2018. A Genomic Imprinting Model of Termite Caste Determination: Not Genetic but Epigenetic Inheritance Influences Offspring Caste Fate. The American Naturalist, 191(6).

Mcafee, A., Pettis, J. S., Tarpy, D. R., and Foster, L. J. 2019. *Feminizer* and *doublesex* knock-outs cause honey bees to switch sexes. PLoS Biology, 17(5): e3000256.

Messerschmidt, D. M., Knowles, B. B., and Solter, D. 2014. DNA methylation dynamics during epigenetic reprogramming in the germline and preimplantation embryos. Genes and Development, 28(8): 812–828.

Mott, B. M., Gadau, J., and Anderson, K. E. 2015. Phylogeography of *Pogonomyrmex barbatus* and *P. rugosus* harvester ants with genetic and environmental caste determination. Ecology and Evolution, 5(14): 2798–2826.

Patalano, S., Vlasova, A., Wyatt, C., Ewels, P., Camara, F., Ferreira, P. G., Asher, C. L., Jurkowski, T. P., Segonds-Pichon, A., Bachman, M., González-Navarrete, I., Minoche, A. E., Krueger, F., Lowy, E., Marcet-Houben, M., Rodriguez-Ales, J. L., Nascimento, F. S., Balasubramanian, S., Gabaldon, T., Tarver, J. E., Andrews, S., Himmelbauer, H., Hughes, W. O. H., Guigó, R., Reik, W., and Sumner, S. 2015. Molecular signatures of plastic phenotypes in two eusocial insect species with simple societies. Proceedings of the National Academy of Sciences, 112(45): 13970–13975.

Pegoraro, M., Marshall, H., Lonsdale, Z. N., and Mallon, E. B. 2017. Do social insects support Haig’s kin theory for the evolution of genomic imprinting? Epigenetics, 12(9): 725–742.

Provataris, P., Meusemann, K., Niehuis, O., Grath, S., and Misof, B. 2018. Signatures of DNA Methylation across Insects Suggest Reduced DNA Methylation Levels in Holometabola. Genome biology and evolution, 10(4): 1185–1197.

Sadd, B. M., Barribeau, S. M., Bloch, G., de Graaf, D. C., Dearden, P., Elsik, C. G., Gadau, J., Grimmelikhuijzen, C. J., Hasselmann, M., Lozier, J. D., Robertson, H. M., Smagghe, G., Stolle, E., Van Vaerenbergh, M., Waterhouse, R. M., Bornberg-Bauer, E., Klasberg, S., Bennett, A. K., Câmara, F., Guigó, R., Hoff, K., Mariotti, M., Munoz-Torres, M., Murphy, T., Santesmasses, D., Amdam, G. V., Beckers, M., Beye, M., Biewer, M., Bitondi, M. M., Blaxter, M. L., Bourke, A. F., Brown, M. J., Buechel, S. D., Cameron, R., Cappelle, K., Carolan, J. C., Christiaens, O., Ciborowski, K. L., Clarke, D. F., Colgan, T. J., Collins, D. H., Cridge, A. G., Dalmay, T., Dreier, S., du Plessis, L., Duncan, E., Erler, S., Evans, J., Falcon, T., Flores, K., Freitas, F. C., Fuchikawa, T., Gempe, T., Hartfelder, K., Hauser, F., Helbing, S., Humann, F. C., Irvine, F., Jermiin, L. S., Johnson, C. E., Johnson, R. M., Jones, A. K., Kadowaki, T., Kidner, J. H., Koch, V., Köhler, A., Kraus, F. B., Lattorff, H. M. G., Leask, M., Lockett, G. A., Mallon, E. B., Antonio, D. S., Marxer, M., Meeus, I., Moritz, R. F., Nair, A., Näpflin, K., Nissen, I., Niu, J., Nunes, F. M., Oakeshott, J. G., Osborne, A., Otte, M., Pinheiro, D. G., Rossié, N., Rueppell, O., Santos, C. G., Schmid-Hempel, R., Schmitt, B. D., Schulte, C., Simões, Z. L., Soares, M. P., Swevers, L., Winnebeck, E. C., Wolschin, F., Yu, N., Zdobnov, E. M., Aqrawi, P. K., Blankenburg, K. P., Coyle, M., Francisco, L., Hernandez, A. G., Holder, M., Hudson, M. E., Jackson, L. R., Jayaseelan, J., Joshi, V., Kovar, C., Lee, S. L., Mata, R., Mathew, T., Newsham, I. F., Ngo, R., Okwuonu, G., Pham, C., Pu, L. L., Saada, N., Santibanez, J., Simmons, D. N., Thornton, R., Venkat, A., Walden, K. K., Wu, Y. Q., Debyser, G., Devreese, B., Asher, C., Blommaert, J., Chipman, A. D., Chittka, L., Fouks, B., Liu, J., O’Neill, M. P., Sumner, S., Puiu, D., Qu, J., Salzberg, S. L., Scherer, S. E., Muzny, D. M., Richards, S., Robinson, G. E., Gibbs, R. A., Schmid-Hempel, P., and Worley, K. C. 2015. The genomes of two key bumblebee species with primitive eusocial organization. Genome Biology, 16(1): 76.

Schneider, C. A., Rasband, W. S., and Eliceiri, K. W. 2012. NIH Image to ImageJ: 25 years of image analysis. Nature Methods, 9(7): 671–675.

Schultz, M. D., Schmitz, R. J., and Ecker, J. R. 2012. ‘Leveling’ the playing field for analyses of single-base resolution DNA methylomes. Trends in Genetics, 28(12): 583–585.

Shaham, R., Ben-Shlomo, R., Motro, U., and Keasar, T. 2016. Genome methylation patterns across castes and generations in a parasitoid wasp. Ecology and Evolution, 6(22): 7943–7953.

Simola, D. F., Graham, R. J., Brady, C. M., Enzmann, B. L., Desplan, C., Ray, A., Zwiebel, L. J., Bonasio, R., Reinberg, D., Liebig, J., and Berger, S. L. 2016. Epigenetic (re)programming of caste-specific behavior in the ant *Camponotus floridanus*. Science, 351(6268).

Singer, M., Kosti, I., Pachter, L., and Mandel-Gutfreund, Y. 2015. A diverse epigenetic landscape at human exons with implication for expression. Nucleic Acids Research, 43(7): 3498–3508.

Standage, D. S., Berens, A. J., Glastad, K. M., Severin, A. J., Brendel, V. P., and Toth, A. L. 2016. Genome, transcriptome, and methylome sequencing of a primitively eusocial wasp reveal a greatly reduced DNA methylation system in a social insect. Molecular Ecology, 25(8): 1769–1784.

Supek, F., Bošnjak, M., Škunca, N., and Šmuc, T. 2011. Revigo summarizes and visualizes long lists of gene ontology terms. PLoS ONE, 6(7).

Vojta, A., Dobrinic, P., Tadic, V., Bockor, L., Korac, P., Julg, B., Klasic, M., and Zoldos, V. 2016. Repurposing the CRISPR-Cas9 system for targeted DNA methylation. Nucleic Acids Research, 44(12): 5615–5628.

Wakefield, R. I., Smith, B. O., Nan, X., Free, A., Soteriou, A., Uhrin, D., Bird, A. P., and Barlow, P. N. 1999. The solution structure of the domain from MeCP2 that binds to methylated DNA. Journal of Molecular Biology, 291(5): 1055–1065.

Wang, X., Wheeler, D., Avery, A., Rago, A., Choi, J. H., Colbourne, J. K., Clark, A. G., and Werren, J. H. 2013. Function and Evolution of DNA Methylation in *Nasonia vitripennis*. PLoS Genetics, 9(10).

Weiner, S. A., Galbraith, D. A., Adams, D. C., Valenzuela, N., Noll, F. B., Grozinger, C. M., and Toth, A. L. 2013. A survey of DNA methylation across social insect species, life stages, and castes reveals abundant and caste-associated methylation in a primitively social wasp. Naturwissenschaften, 100(8): 795–799.

Winkler, A. M., Webster, M. A., Vidaurre, D., Nichols, T. E., and Smith, S. M. 2015. Multi-level block permutation. NeuroImage, 123: 253–268.

Woodard, S. H., Lozier, J. D., Goulson, D., Williams, P. H., Strange, J. P., and Jha, S. 2015. Molecular tools and bumble bees: Revealing hidden details of ecology and evolution in a model system. Molecular Ecology, 24(12): 2916–2936.

Zilberman, D. 2017. An evolutionary case for functional gene body methylation in plants and animals. Genome Biology, 18(1): 17–19.

