## Supplementary 2 for "Methylation and Gene Expression Differences Between Reproductive Castes of Bumblebee Workers"

1) Department of Genetics and Genome Biology
University of Leicester
Leicester

### 2.0: Permutation Analysis

Permutation tests are used to randomly shuffle labels across a given data set to ensure results obtained via a significance test are due to biological causes rather than random variation within the data resulting in type I errors.

Here we employed a similar method to Arsenault et al. (2018) and Libbrecht et al. (2016). After removal of positions containing zero methylation for every sample and filtering by coverage, the sample labels for data at each CpG were randomly shuffled 10,000 times. A logistic regression was carried out for each random data set using methylKit. The number of differentially methylated sites per permutation were plotted (Fig.S1).

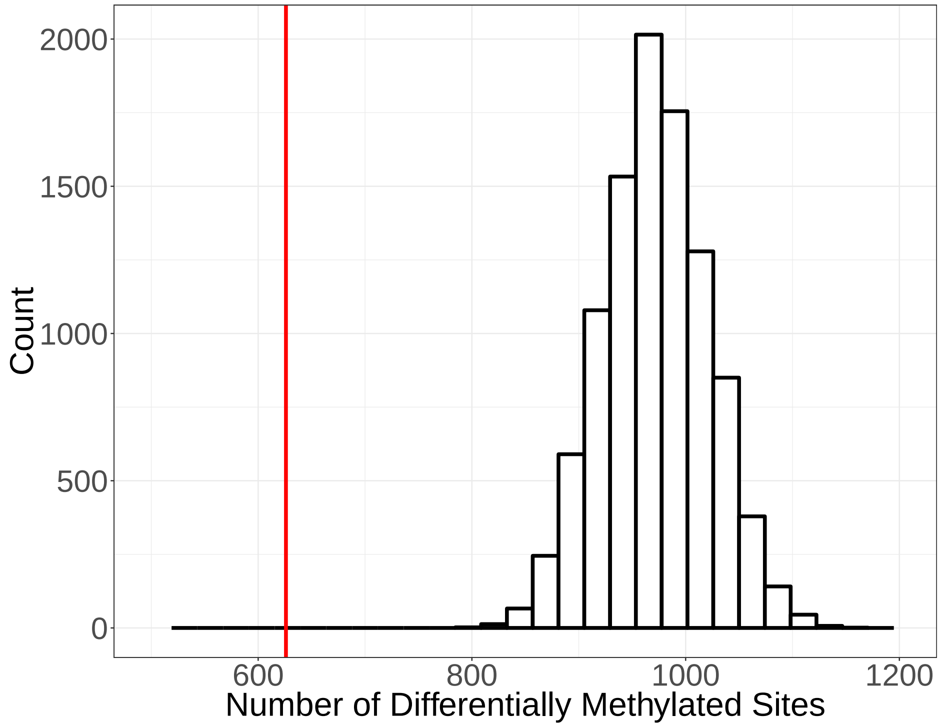


Figure S1: Histogram of the number of differentially methylated sites obtained from 10,000 permutations. The red line indicates the number of sites observed from the un-shuffled data.

More sites are found to be differentially methylated using the random data as variation between colonies is higher than variation caused by reproductive status (Fig.S2a and S2b). Using this method 'colony' can no longer be taken into account as a consistent covariate and the effect therefore inflates the number of obtained differentially methylated sites.

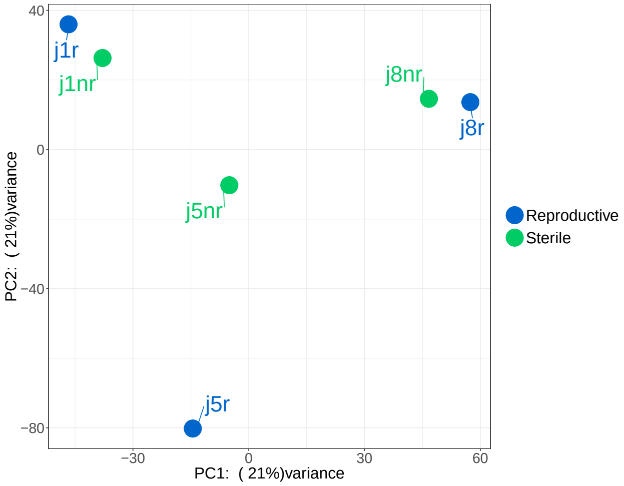


(a) PCA Plot

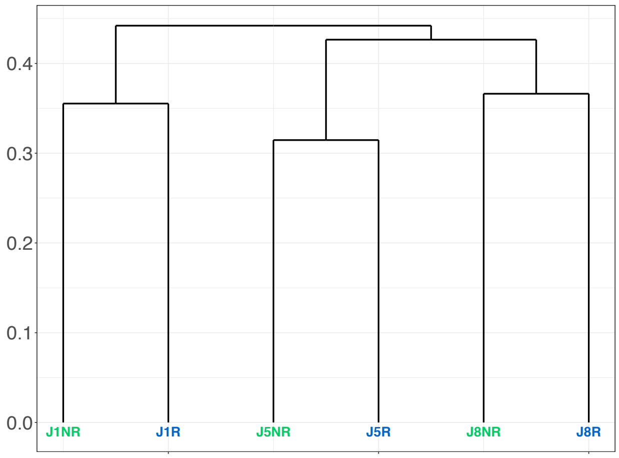


(b) Dendogram

Figure S2: (a) PCA plot showing samples cluster more closely by colony than by reproductive status. (b) Dendogram showing sample cluster by colony. Red labels indicate reproductive samples and blue labels represent sterile samples.

Whilst permutation tests are useful for some data sets, when structure is present in the data they become unreliable, as discussed in Winkler et al. (2015). A higher number of replicates would allow label shuffling within confounding factors, maintaining the structure of the data, thus allowing a valid permutation.

### 2.1: Additional Methylation Analyses

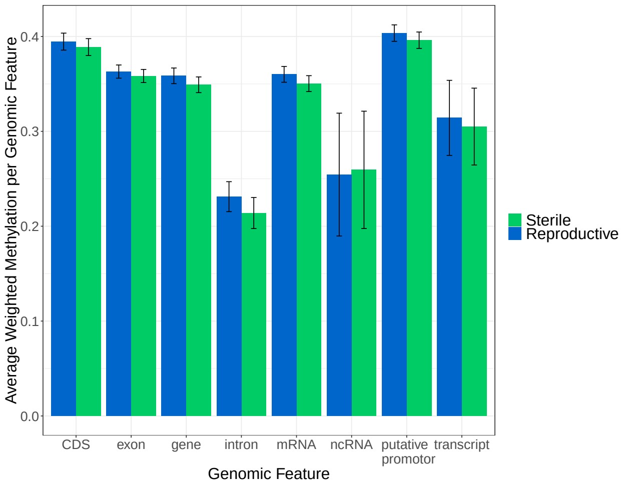


(a)

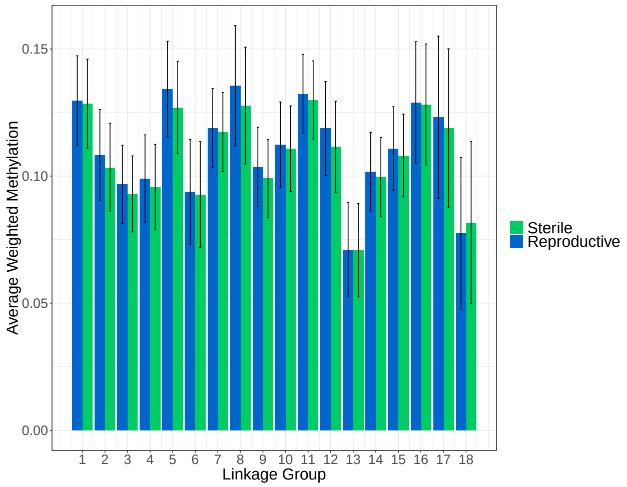


(b)

Figure S3: (a) The mean weighted methylation level across colonies for each genomic feature for both reproductive and sterile workers. Error bars are 95% confidence intervals of the mean. This graph includes a putative promotor region, this was defined as 5000bp upstream of each gene annotation. As the current annotation file for *B. terrestris* does not include promotor regions. This result should be viewed with care as some of the 'putative promtor' regions also contain other annotated genomic features such as other genes and coding regions. (b) The mean weighted methylation level across colonies for each linkage group defined in the B. terrestris genome Bter_1.0 (NCBI Accession: GCA_000214255.1). Error bars are 95% confidence intervals of the man. This figure should also be viewed with caution as currently there are >5000 unplaced scaffolds for this genome build which are not included in the figure.

### 2.2: Sample Exclusion from Differential Expression

Sample J8_24 was classed as reproductive but clustered with the sterile samples in both a principle component analysis (PCA) and a poisson distance matrix, taking into account expression levels of all genes (Fig.S4a and S4b). It also clustered with sterile samples in a hierarchical cluster using euclidean distances based on the top 100 differentially expressed genes (Fig.S4c). Including this sample led to a decrease in the number of differentially expressed genes identified (110 with sample, 334 without sample). After removal of this sample all other samples clustered by reproductive status (Fig.S5a, S5b and S5c).

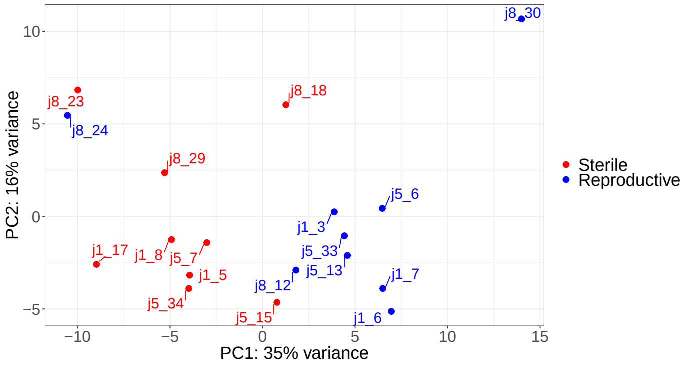


(a) PCA Plot

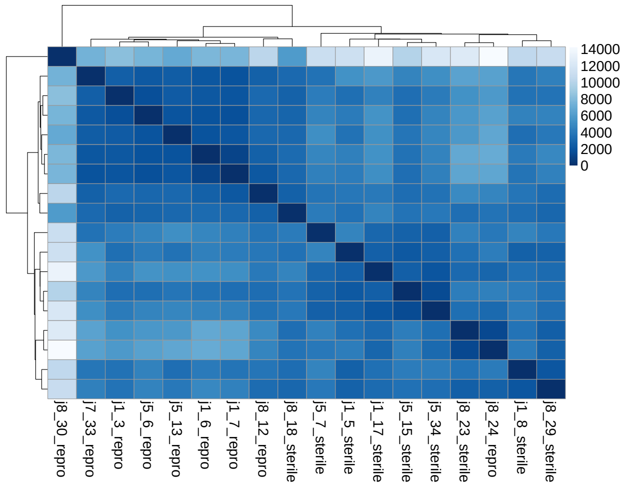


(b) Poisson Distance Matrix

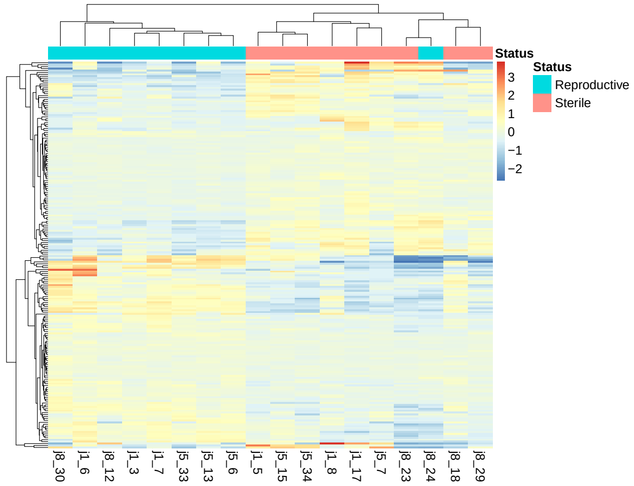


(c) Elucidean Cluster Heatmap

Figure S4: Graphs generated from differential expression analysis using DESEQ2 for all samples. (a) PCA plot showing the different reproductive status' grouped together. (b) Poisson measurement of dissimilarity between counts. (c) Heatmap showing the top 100 differentially expressed genes in reproductive and sterile workers. Red colour indicates over-expression and blue colour indicates under-expression.

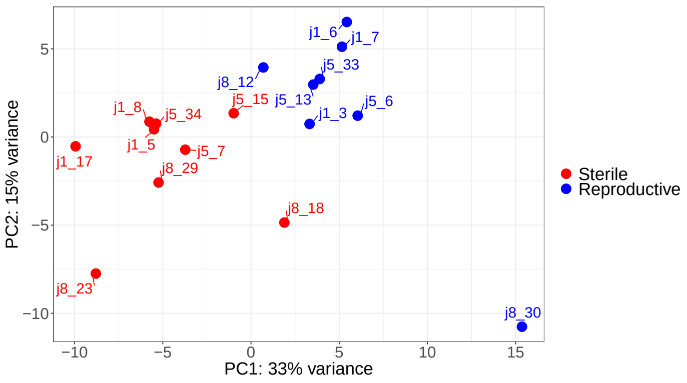


(a) PCA Plot

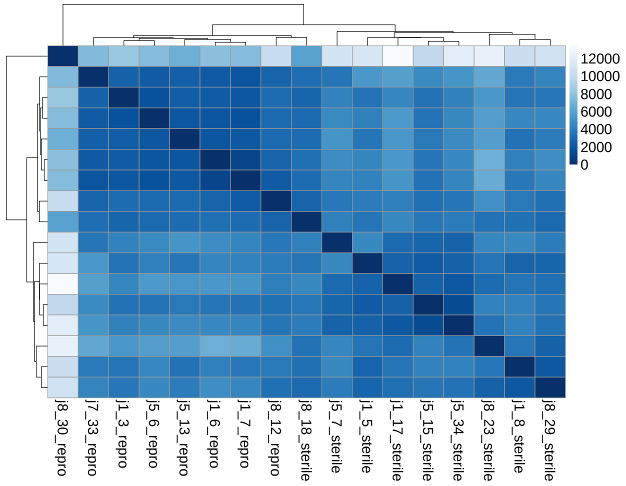


(b) Poisson Distance Matrix

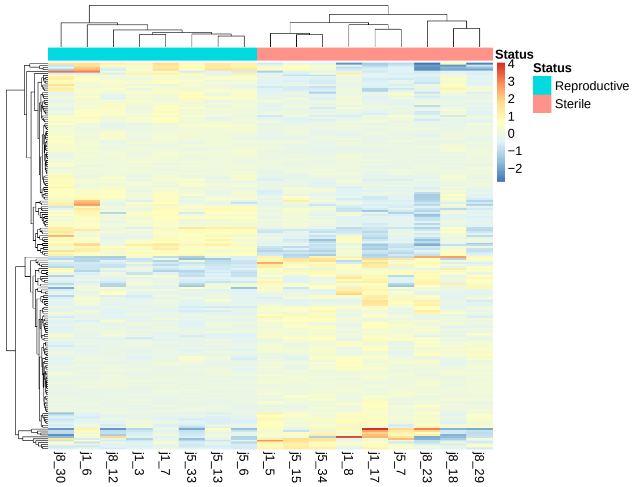


(c) Elucidean Cluster Heatmap

Figure S5: Graphs generated from differential expression analysis using DESEQ2 for all samples, excluding J8_24. (a) PCA plot showing the different reproductive status' grouped together. (b) Poisson measurement of dissimilarity between counts. (c) Heatmap showing the top 100 differentially expressed genes in reproductive and sterile workers. Red colour indicates over-expression and blue colour indicates under-expression.

### References


Samuel V. Arsenault, Brendan G. Hunt, and Sandra M. Rehan. The effect of maternal care on gene expression and DNA methylation in a subsocial bee. *Nature Communications*, 9(3468), 2018. ISSN 20411723. doi: 10.1038/s41467-018-05903-0. URL http://dx.doi.org/10.1038/s41467-018-05903-0.

Romain Libbrecht, Peter Robert Oxley, Laurent Keller, and Daniel Jan Christoph Kronauer. Robust DNA Methylation in the Clonal Raider Ant Brain. *Current Biology*, 26:1–5, 2016. ISSN 09609822. doi: 10.1016/j.cub.2015.12.040. URL http://linkinghub.elsevier.com/retrieve/pii/S0960982215015717.

Anderson M. Winkler, Matthew A. Webster, Diego Vidaurre, Thomas E. Nichols, and Stephen M. Smith. Multi-level block permutation. *NeuroImage*, 123:253–268, 2015. ISSN 10959572. doi: 10.1016/j.neuroimage.2015.05.092. URL http://dx.doi.org/10.1016/j.neuroimage.2015.05.092.
